# Precision localization of cellular proteins with fluorescent Fab-based probes

**DOI:** 10.1101/2021.10.18.464886

**Authors:** F. Liccardo, M. Lo Monte, B. Corrado, M. Veneruso, S. Celentano, M.R. Coscia, G. Coppola, P. Pucci, G. Palumbo, A. Luini, M. Lampe, V.M. Marzullo

**Author notes:** Current address: Cardiovascular Research Institute, University of California San Francisco, San Francisco, USA. Equal contribution (F. Liccardo, M. Lo Monte).

## Abstract

Currently, a major technical limitation of microscopy based image analysis is linkage error – a visualization error that is measured by the distance between the cellular protein to be detected and the fluorescence emitter used for detection. With continuously improving resolution of today’s (super-resolution) microscopes, the linkage errors can severely hamper the correct interpretation of images and is usually introduced in experiments by the use of standard intracellular staining reagents such as fluorescently labelled antibodies. The linkage error of standard labelled antibodies is caused by the size of the antibody and the random distribution of fluorescent emitters on the antibody surface. Together, these two factors account for a fluorescence displacement of ~40nm when staining proteins by indirect immunofluorescence; and ~20nm when staining with fluorescently coupled primary antibodies. In this study, we describe a class of staining reagents that effectively reduce the linkage error by more than five-fold when compared to conventional staining techniques. These reagents, called Fluo-N-Fabs, consist of an antigen binding fragment of a full-length antibody (Fab/ fragment antigen binding) that is selectively conjugated at the N-terminal amino group with fluorescent organic molecules, thereby reducing the distance between the fluorescent emitter and the protein target of the analysis. Fluo-N-Fabs also exhibit the capability to penetrate tissues and highly crowded cell compartments, thus allowing for the efficient detection of cellular epitopes of interest in a wide range of fixed samples. We believe this class of reagents realize an unmet need in cell biological super resolution imaging studies where the precise localization of the target of interest is crucial for the understanding of complex biological phenomena.

## INTRODUCTION

Far-field fluorescence microscopy is constantly improving in the continuous attempt to resolve the finest details of intracellular architectures by overcoming the Abbe limitation^1^. This limitation, known as the diffraction limit, prevents the resolution of structures smaller than approximately half the wavelength of light and, as a consequence, makes it impossible to resolve, with high precision, cellular nanostructures that are <250 nm in size. To overcome this limit, super resolution microscopy techniques have been developed. The most relevant improvements to super resolution methodologies consist of a combination of single molecule localization methods (SMLMs), like STORM, and the deterministic approach, that have led to an increased resolving power of microscopes up to a very few nanometres ^2^ ^3^ ^4^ ^5^. This unprecedented resolving power of modern instruments ^6^ ^7^, combined with recently developed algorithms for image processing ^8^ ^9^ ^10^ and for the evaluation of the resolution limit 11 have modified the resolution problem in diffraction-unlimited imaging.

Nevertheless, despite the technological advancement of super resolution microscopy, the localization of precise coordinates for a given cellular protein is still a challenging task. This is essentially due to the fact that in all the fluorescence microscopy techniques, the preferred fluorescence emitters are organic molecules conjugated to probes that bind directly to the cellular epitope of interest. The use of fluorescently labelled probes introduces an error in the localization of the object of interest as the actual position of the biological target within a cell differs from the experimentally determined location due to the distance between the fluorescence emitter and the target epitope ^12^. The localization uncertainty introduced by the presence of a linker that physically ties the cellular target with the emitters is called linkage error ^13^. This represents a major problem in super resolution imaging analyses since an error with an order of magnitude greater than the nanostructures being visualized can introduce distortions in their reconstruction, especially when performing SMLM studies. In SMLMs, linkage errors are identified as the space between the fluorescent label and the target protein; thus, large probes and the distributions of the fluorescence emitters on the probes influence the magnitude of the linkage error ^14^ ^15^. These limitations are of particular concern when utilizing the largest and most extensive class of detection probes used as microscopy reagents – fluorescently labelled immunoglobulins/antibodies. These use of these reagents in super resolution microscopy is characterized by a target localization uncertainty due mainly to two reasons: the size of the antibody (≈12 nm ^16^ along the longest axis) and the random distribution of emitters on the antibody probe surface due to the uncontrolled chemical conjugation reactions used to synthesize these probes.

So far, several studies have shown that indirect (with labelled secondary antibodies) and direct (with labelled primary antibodies) immunofluorescence techniques can produce a fluorescence displacement of the labels from the cellular target of interest by about 40 nm and 20 nm, respectively ^12^ ^14^ ^17^. When this displacement is measured on the surface of a sphere centred on the fluorophore, the experimental location of the target protein is calculated to be at a distance of up to 20 nm away from the actual intracellular location [see **Figure 1A** for a pictographic representation of this concept].

**Figure 1:**
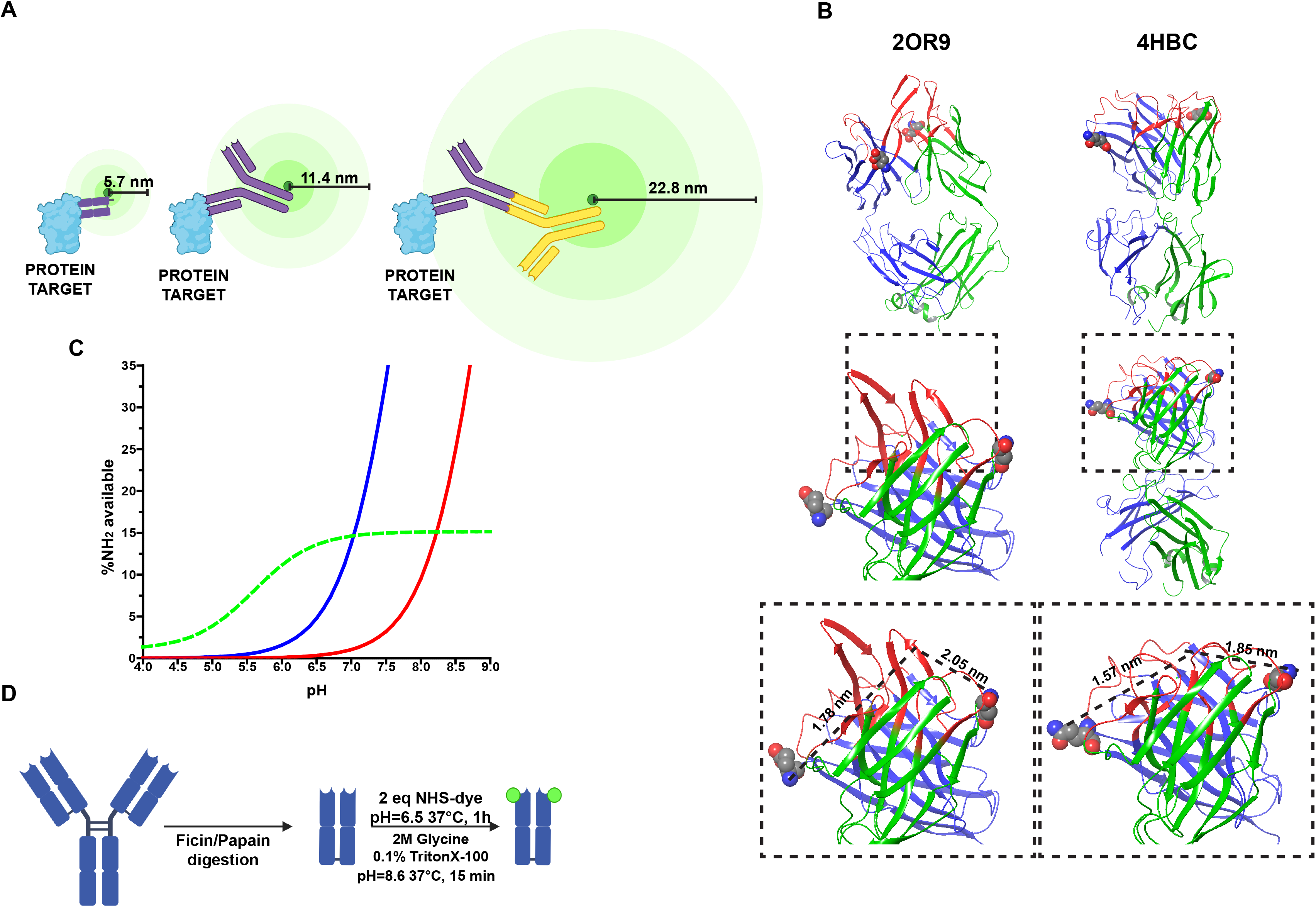
Linkage error and probes rational design. **A)** Representation of the fluorescent displacement from the protein target produced from secondary (yellow), primary (purple) antibodies and Fab fragment (purple) labelled on the most remote area of their respective structures **B)** Crystal structures of the mouse monoclonal Fab 2OR9 and the rabbit monoclonal 4HBC. The light chains are showed in green; heavy chains are showed in blue; the epitope binding zone of both the Fab fragment is highlighted in red and the N-terminal amino-groups of both the heavy and light chains of each Fab are represented with spheres. The relative distance between the N-terminal amino groups and the epitope binding zone are in the range of 1.5-2 nm independently from the different antibody sources **C)** Percentage of free amino groups at different pH. In blue is shown the percentage ratios of non-protonated N-terminal α-amino group of the 2OR9 Fab heavy chain (pKa=7.8) in a range of pH between 4 and 9; in red there is the percentage of non-protonated ε-amino group of a lysine (pKa= 8.98) in a range of pH between 4 and 9; in green is shown the ratio between the percentage ratio of not protonate N-terminal α-amino group and not protonated ε-amino group of the lysine. **D)** Fluo-N-Fab reaction scheme.

Moreover, due to the nature of random labelling, primary antibodies conjugated to fluorescence markers are potentially labelled in areas far removed from the epitope binding zone, such as the C-terminal region of the Fc fragment. This would imply an estimated distribution of the labelling in a fluorescence sphere with a radius of 11.4 nm during the use of full-length primary antibodies as probes and 22.8 nm during the use of labelled secondary antibodies. This results in a corresponding fluorescence displacement of 22.8 nm and 45.6 nm respectively, resulting in a mislocalization of intracellular target coordinates (**Figure 1A**).

The most obvious and immediate method for the reduction of linkage error is the development of probes and detection methods alternative to the large full-length antibodies 18 19. Here we do not consider the potential alternatives represented by fluorescent tags such as photo-switchable fluorescent proteins, that have found applications in SMLM techniques like PALM ^20^. Indeed, CRISPR technologies allow for the labelling of endogenous genes such that protein abundance is not altered and all target proteins are labelled. However, fluorescent proteins are ~2–5 nm in size and are attached to the protein of interest by means of an amino acid linker of a length of up to ~5 nm. Hence, the imaged location of a protein deviates from the true location by the vector sum of these quantities ^21^, resulting in a significant linkage error. Moreover, the presence of genetically engineered tags can potentially perturb the function and location of the target protein ^22^. Fluorescent proteins are also characterized by low photon output and blinking features ^23^ ^24^ which may hamper SMLM detection. Finally, and most importantly, genetically encoded fluorescent proteins cannot be easily utilized for the application of SMLM techniques in primary cell lines or biopsies.

What is a viable strategy then to reduce the linkage error in microscopy-based imaging studies? One way is by using significantly smaller probes in order to decrease the distance between the target of interest and the fluorescent label ^18^. Some of these probes are non-antibody based chemicals such as DARPins ^25^ and affimers ^26^. So far, despite the small size of these reagents (~2 nm), DARPins have not been well explored for their potential use in super resolution imaging, while affimers have found limited success when applied to STORM microscopy experiments ^27^ ^28^. Between all the alternative probes used, the most efficient are the nanobodies ^29^ ^30^. This class of immunoglobulins are defined as single-domain variable fragments of camelid-derived heavy-chain antibodies. Nanobodies exhibit useful properties such as a small size (2.5-4 nm) ^13^ ^31^ ^32^ ^33^. However, the main limitation for all these alternatives to the antibodies is that they are not available for a vast majority of protein targets and require considerable time and effort for their preparation. These aspects have limited the frequency and routine use of these reagents in biology laboratories.

Here, we present a class of reagents called Fluo-N-Fabs (Fab-Fluorophore conjugates) that precisely label endogenous proteins as visualized by the SMLM STORM. Specifically, this class of precision labelling reagents are based on the use of antigen binding fragments (Fabs) obtained by the enzymatic digestion of commercially available primary antibodies. We have developed a versatile and reproducible protocol for the selective conjugation of the N-terminal amino group of the Fab fragments. The use of Fluo-N-Fabs as reagents for super resolution microscopy shows multiple advantages. The Fab is a small portion of IgG antibodies (4-5 nm) that successfully identify the cellular target of interest via the epitope binding zone. We selectively labelled the N-terminal amino group of the Fab heavy and light chains as these are located in close proximity (within 2 nm) of the epitope-binding zone. The distance between the N-terminal amino groups and the epitope binding zone is highly conserved in different IgGs isotypes and species ^34^. The small size of the Fab and position of the dye at the N-terminal amino groups of the Fab locates the emitters very close to the surface of the protein of interest.

Another important advantage in the application of Fluo-N-Fabs is their capability to penetrate crowded cell compartments in fixed cell samples and potentially in biopsy tissues (i.e. formalin-fixed paraffin-embedded tissues) ^35^ ^36^ ^37^ ^38^. This property is intrinsically related to the relative small size of the Fab fragment and can be conveniently exploited in combination with the N-terminal site specific conjugation protocol to obtain a stoichiometric marking of cellular epitopes. A control of the size of the linker and the number / position of the fluorophores with which it is conjugated represents a considerable advantage for quantitative imaging using super resolution microscopy ^39^ ^40^ ^41^ ^13^ ^42^.

Finally, other advantages in the use of Fluo-N-Fabs is their availability, speed of production and accessibility for any cell biology laboratory. Fab fragments can be easily obtained by enzymatic digestion of potentially every full-length primary antibody ^43^ ^44^. Given the enormous interest in the development of primary full-length antibodies for various cell biology applications, potentially vast repertoires of antibodies targeting the most diverse cellular epitopes are commercially available that can be exploited for the generation of our precision labelling reagents.

In sum here we present a promising class of imaging reagents that are easy to produce in the lab with a simple chemical protocol. These reagents reduce the linkage error in single molecule localization microscopy techniques, like STORM, and allow the precision labelling of cellular nanostructures.

## RESULTS

### REGIOSELECTIVE Fab MODIFICATION THROUGH pH MEDIATED ACYLATION

Different site-selective chemical modifications of proteins have been developed exploiting the reactivity between a target amino acid and a specific functional group exposed by the molecule to be conjugated (such as a fluorescent dye) ^45^. Protein residues that are susceptible to modifications are primary amino groups on the N-terminal most amino acid (the first amino acid on a peptide chain) or on lysine side chains, thiols (cysteine side chains), C-terminal residues (carboxylic acid, glutamic acid) and hydroxyl groups. Of these residues, N-terminal amino groups and lysine side chains are the most commonly functionalized for modifications. Typically, the N-terminal amino groups are solvent exposed and, being readily accessible ^46^, are ideal targets for various site-selective modification reactions.

To label the N-terminal amino groups, N-Hydroxysuccinimide (NHS) esters (or their more soluble sulfo-NHS analogues) are the most efficient reactive species that are inserted into fluorophores to conjugate proteins ^47^ ^48^. The fluorophores containing NHS esters then readily acylate primary amino groups on proteins, yielding a stable amide bond.

Acylation through the use of NHS esters has practical advantages - like mild reaction conditions and high stability of the final conjugated products ^49^. However, NHS esters do not discriminate between reacting with amino groups on either the N-terminal amino acid or amino groups on lysines ^50^. To overcome this limitation, it is in principle possible to induce reaction selectivity using NHS esters by strictly controlling the reaction pH conditions since the pKa value of the lysine side chains are comparably high, with average of 7.7 ± 0.5 ^51^. Therefore, the acylation of primary amino groups, performed with controlled pH reaction conditions, may provide sufficient regioselectivity favouring the N-terminal coupling ^52^. By exploiting this difference in pKa values, we attempted to achieve a selective acylation of N-terminal amino groups in Fab fragments generated after the enzymatic digestion of primary antibodies.

Towards this purpose, we first undertook an *in silico* approach to examine the theoretical pKa of the N-terminal amino group and the distance between the epitope binding zone and the N-terminal amino group on Fab fragments whose crystal structures have been solved and whose molecular architecture represents the two classes of antibodies that are commercially available and of wide use – namely an IgG monoclonal rabbit Fab (PDB: 4HBC) and an IgG_1_ monoclonal mouse Fab (PDB: 2OR9) (**Figure 1B)**. Both the Fab fragments were analysed using a hybrid approach by first applying a focused quantum-mechanical strategy that was superimposed on a more common molecular-mechanical environment (**see Supplementary materials**). This approach yielded a 3D-structural characterization of the Fab fragments that were now amenable to computational prediction of pKa values of both the N-terminal amino group and the amino group of the lysine side chain. **Table 1** reports the pKa values of the amino groups of the two Fabs as obtained by *in silico* calculations. For both Fab fragments, we determined a high mean pKa of 11 – 12 for 90% of the lysine side chain amino groups, while the remaining 10% had a pKa value of ~9. These results are in line with previously published studies ^53^ ^54^. Surprisingly, the pKa value of the N-terminal amino group for the antibodies derived from different species was significantly different. For the Fab fragment derived from rabbit, the calculated pKa for both the heavy and light chains was ~7.5, while for the mouse Fab fragment, the pKa for the light chain was ~9.96 and for the heavy chain was ~7.8. This difference can be explained by the position of the N-terminal amino group in the crystal of the mouse Fab. This N-terminal amino group on the light chain interacts with two glutamic side chains at positions 27 and 93 of the light chain (**Supplementary Figure 1A**), thus stabilizing the protonated form with salt bridges. This coordination can result in an unusually high pKa value for this particular N-terminal amine ^51^ ^55^. To support these results, we further analysed 124 amino acid sequences of the variable domain of IgG1 mouse k isotype antibody sequences that are available in the International Immunogenetics information system database, in order to understand the frequency of N-terminal amino group coupling to internal glutamic acid residues. We discovered that while the glutamic acid 27 was highly conserved, glutamic acid 93 exhibited poor conservation (**Supplementary Figure 1B**), suggesting that the particular mouse Fab fragment utilized for the *in silico* analyses was unique. We hence expanded the repertoire of crystal structures for IgG1 monoclonal mouse Fab fragments and analysed 16 different structures to determine the pKa of their respective N-terminal amino groups. We observed that only two of them had pKa values similar to the one previously analyzed (2OR9) (**Table 2**). This analysis determined that in a majority of the Fab’s considered, the pKa values of the two chains show a mean value of 7.5. However, in a small number of Fab fragments, one of the two amino terminal groups can have a high pKa value. Our *in silico* analyses suggested that in a majority of Fab fragments, a high degree of confidence is attributed to the probability that at least one N-terminal amino group is available for chemical modification with NHS-ester containing fluorophores at neutral pH.

**Table 1:**
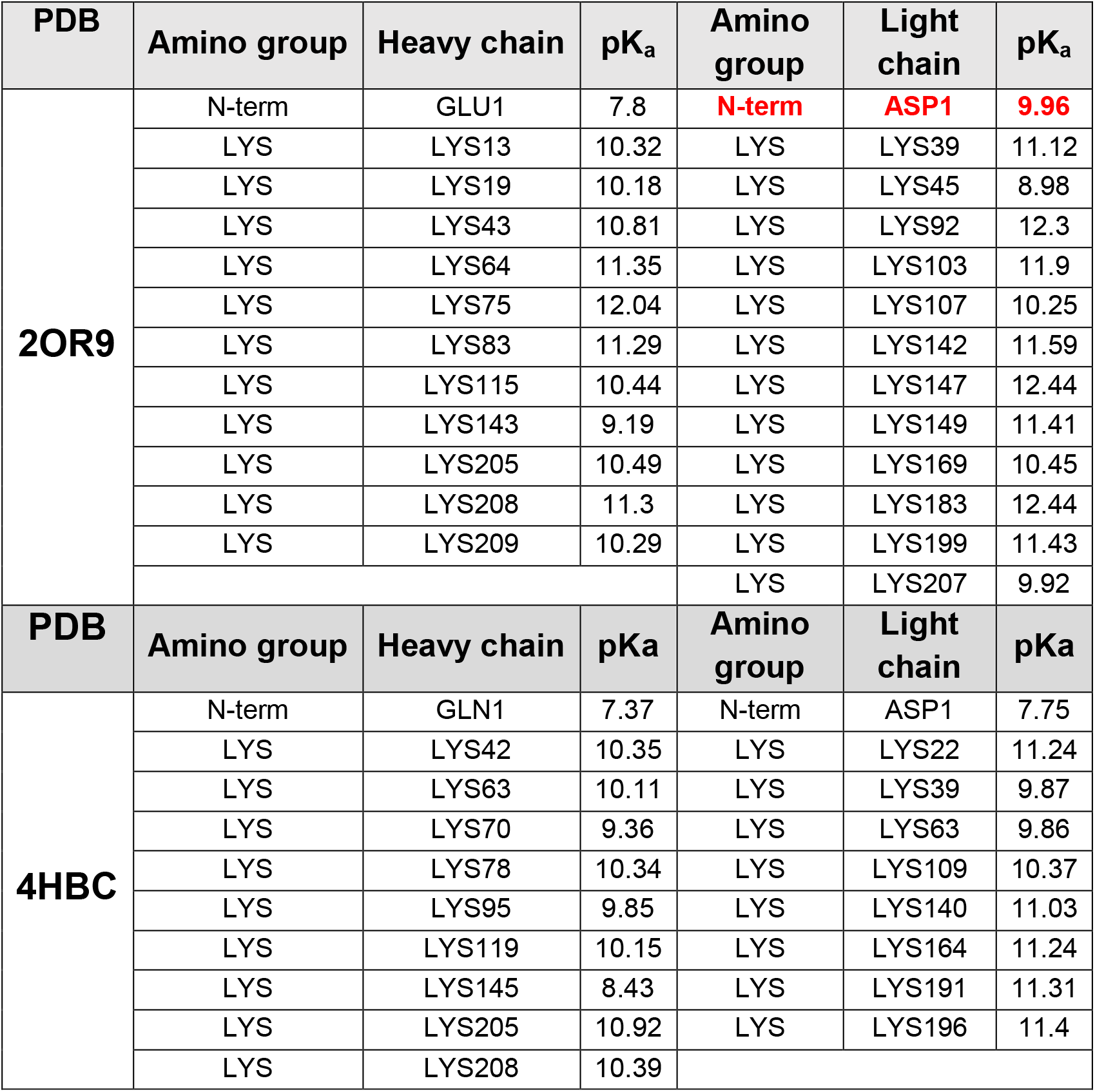
predicted pKa values of heavy and light chains Fabs amino groups.

**Table 2:** Prediction of pKa values of heavy and light chain N-terminal amino groups of mouse monoclonal Fab fragments

| PDB code | N-terminal Light chain pKa | N-terminal Heavy chain pKa |
| --- | --- | --- |
| 1CBV | 7.49 | 8.02 |
| 1BAF | 7.64 | 7.83 |
| 5KVD | 7.64 | 7.55 |
| 3CFC | 8.76 | 7.87 |
| 4PB9 | 7.83 | 8.05 |
| 1H0D | 8.05 | 8.17 |
| 1EJO | 9.11 | 7.45 |
| 1D6V | 8.1 | 7.84 |
| 3L5W | 8.56 | 7.84 |
| 4L5F | 7.98 | 8.23 |
| 2ADF | 6.39 | 7.84 |
| 3HZK | 7.7 | 7.98 |

Taking these values into account we have made a series of simulations comparing the reactivity of an N-terminal group with pKa to 7.8 with that of a lysisine side chain amino groups with pKa 8.98. Specifically, we simulated a pH titration curve and evaluated the percentage ratios of non-protonated individual amino groups (NH_2_) by calculating the ratio of NH_2_/NH_3_+ between the N-terminal amino group and the ε-amino group of the lysine. Our results indicated that an increase in the ratio reduced the pH of the conjugation reaction (i.e.) a decrease of 0.5 pH units implies an increased selectivity for the labelling occurring on the N-terminal amino group as the NH_2_/NH_3_+ ratio increases by a factor >5. We then estimated that the percentage of non-protonated amino groups available at a range of pH reaction conditions (**Figure 1C**) to determine which pH corresponds to a maximum percentage of selectivity achievable in our reaction. We determined that at a pH = 6.5, achieved a reaction selectivity toward the N-terminal amino group of 90%. We hence selected the pH = 6.5 as the preferred pH for the conjugation reaction considering that lysines with a pKa value in that range are very rare (see above) ^53^ ^54^ (**Figure 1C and 1D**). Under these conditions, based on the above considerations, the lysine conjugation is extremely rare.

Once we established the criterion of pH selection, we developed a strategy to label Fab fragments with an NHS-ester coupled fluorescent dye (**Supplementary Fig 2** and Methods). Upon completion of the reaction, the products were subjected to quality control to determine the selectivity of labelling product and degree of labelling by high pressure liquid chromatography (HPLC) analyses (**Supplementary Fig 3**). To test our reaction protocol, we first generated Fabs by enzymatic digestion of two full-length monoclonal antibodies that are routinely used in cell biology laboratories – one that recognizes and binds to the ten amino acid peptide tag EQKLISEEDL/Myc (Fab_Myc_) and another that recognized and binds to the cytoskeletal protein tubulin (Fab_tubulin_). The digested Fab fragments were then incubated with a NHS ester CF568 fluorescent dye at a pH = 6.5 (see Methods). To separate the individual Fab fragments conjugated with varying amounts of CF568, the reaction products were purified by HPLC using size exclusion chromatography ^44^ and hydrophobic interaction chromatography (HIC) ^56^.

The HIC chromatogram of both the Fab fragments suggested a high level of selectivity of the reaction. Indeed, the chromatogram of Fab_tubulin_ labelled with the CF 568 is characterized by the presence of only two fluorescent populations (**Supplementary Fig 3A**) compatibly with the availability of two non-protonated N-terminal residues available for the reaction that can produce a distribution of monolabelled and dilabelled products. Similarly, the HIC chromatogram of the Fab_Myc_ labelled with CF 568 analyses showed the presence of only a single labelled product (**Supplementary Fig 3B**) as expected since the high pKa value of the light chain N-terminal amino group. This narrow distribution of fluorescent peaks suggested a site selective labelling of the Fab fragments at the N-terminal amino group, since a random labelling on lysines would lead to a widely heterogeneous distribution of fluorescently labelled products ^57^. To confirm this reaction selectivity, we enriched the labelled Fab_Myc_ fractions by HIC and sequenced them by peptide mapping and mass spectrometry. We observed that among all the peptides that were sequenced, only one peptide – EVHLVESGGDLVKPGGSLK was positively labelled with CF 568 (**Supplementary Fig 4**). An analysis of the crystal structure of the full length Myc antibody (PDB: 2OR9) confirmed that this labelled peptide mapped to the N-terminal region of the heavy chain. Importantly, we did not observe other CF 568 carrying peptides, further supporting the notion that at a reaction pH = 6.5 all the other amino groups (i.e. lysines) should be fully unavailable for conjugation with fluorophores.

In summary, the HIC chromatography and peptide mapping analyses indicated the presence of a single labelled peptide on the heavy chain of the Fab fragments. Our results also indicated that the Fab light chain does not to carry any detectable labelled peptides, as expected from our *in silico* analyses of the reaction pKa.

### IDENTIFICATION OF FLUOROPHORES SUITABLE FOR THE N-TERMINAL LABELLING

The selective labelling of the N-terminal group of the Fab fragment involves the introduction of a substantial modification on a residue that is in close proximity with the epitope binding zone. This modification might potentially impact the affinity or penetration property of the final probe. Our success in conjugating a CF 568 fluorophore to the Fab_Myc_ and Fab_tubulin_ fragments prompted us to determine the repertoire of commercially available fluorescent dyes that are A.) available for conjugation under our established reaction conditions and B.) preserve the affinity for the epitope of the N-terminally conjugated Fab.

To address this, we selected a subset of nine commercially available fluorophores that found many applications in super resolution microscopy. The fluorophores selected are: AF 647, AF 532, ATTO 647N, ATTO 647, ATTO 590, ATTO 565, Star 635P, Star 635 and Star Red.

We conjugated the selected fluorophores to the Fab_tubulin_ fragment. Importantly, each independent conjugation reaction yielded Fluo-N-Fabs with similar DOLs. We then evaluated the capability of the obtained probes to bind the tubulin in fixed cells first by imaging of the tubulin staining in HeLa cells using confocal fluorescent microscopy, and, secondly by assessing the dissociation rates of each one of the Fab_tubulin_ Fluo-N-Fab as relative measure of their binding affinity.

The imaging of tubulin staining in HeLa cells showed that the Fabs conjugated with ATTO (ATTO 565, ATTO 590, ATTO 647 and ATTO 647N) dyes display a near complete loss of affinity of the Fab fragment towards the antigen (**Figure 2A, bottom panel**), with the appearance of unspecific fluorescent nuclear aggregates that resembled staining patterns obtained with randomly labelled nanobodies ^33^). We then determined the capacity of all other Fluo-N-Fabs to efficiently stain tubulin (**Figure 2A**). The various conjugates efficiently stained endogenous tubulin with the appearance of filamentous substructures within HeLa cells (**Figure 2A, panel 1**). The images were comparable to the staining patterns obtained using either conjugated full-length primary antibodies or secondary antibodies. We hence excluded the Fab_tubulin_ - ATTO conjugates from further analyses of binding affinity

**Figure 2:**
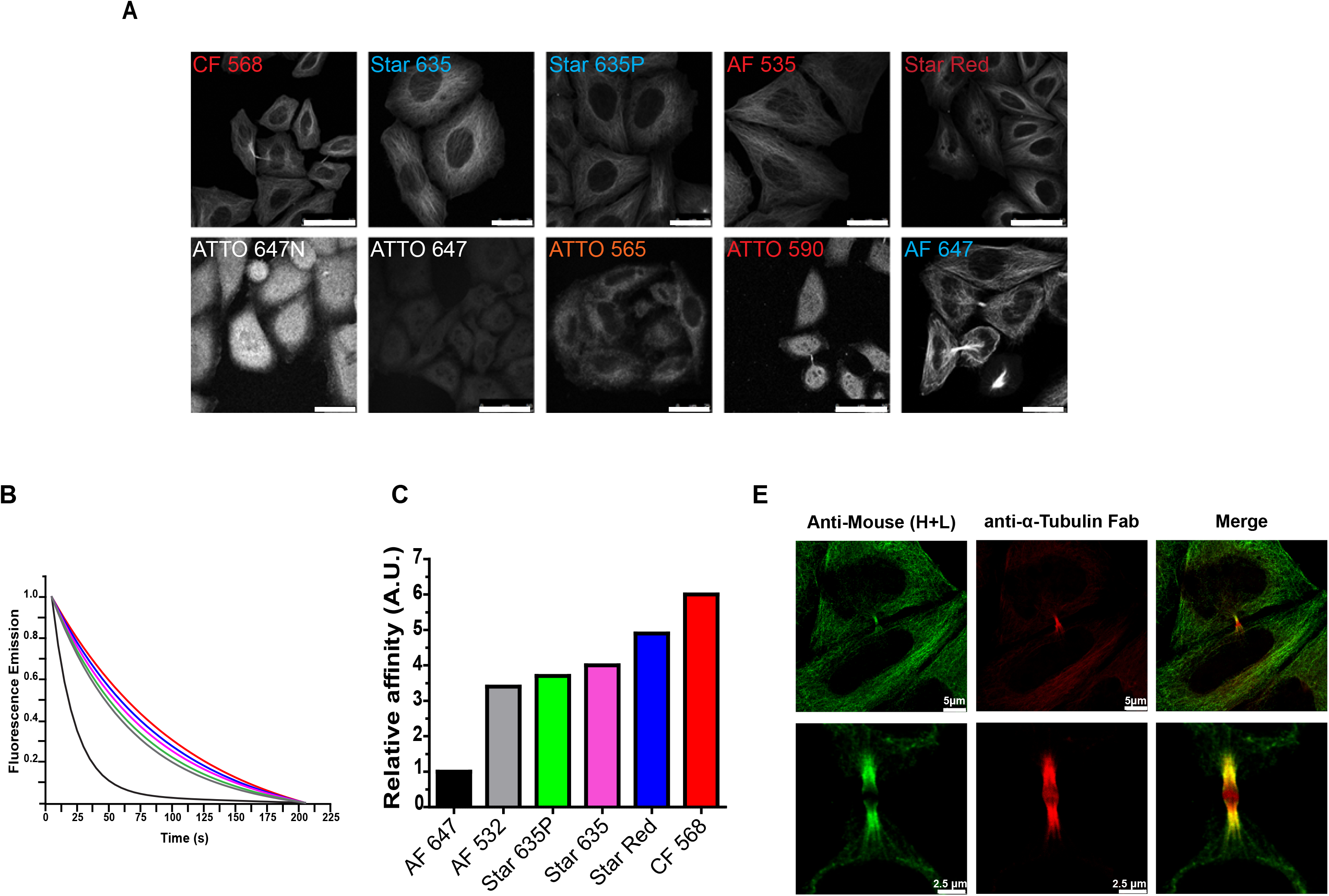
Fluorophore structural properties modulate the Fluo-N-Fabs affinity. **A)** Tubulin staining of Hela M cells obtained using anti-α-tubulin Fab labelled with different type of fluorophores. From the left to the right: CF 568, Star 635, Star 635P, AF 535, Star Red, ATTO 647N, ATTO 647, ATTO 565, ATTO 590 and AF 647 (Scale bar: 50 μm) **B)** Normalized fluorescence emission of each probe vs Time: black lane AF 647 probe, red lane CF 568 probe, blue lane Starred probe, pink lane Star 635 probe, green lane Star 635P probe, grey lane AF 532 probe **C)** relative affinity of each probe for tubulin obtained considering the ratio T_1_ (AF 647 probe)/T_1_ (probe), where T_1_ is the fast decay component of the bi-exponential equation used to fit the dissociation curves of each probe. **D)** Midbody staining of HeLa cells using anti-α-tubulin Fab labelled with CF 568 (red) as primary binders and anti-Mouse Alexa Fluor 488 as secondary antibody (green).

To further understand the different properties of the tested Fluo-N-Fabs, we analysed a set of critical parameters for fluorophore selection under our reaction conditions by an *in-silico* analysis of the physico-chemical properties of the ten fluorophores used in this study.

We initially calculated a set of 90 descriptors and progressively narrowed them by applying statistical methods to obtain a curated set of five parameters (Molecular weight, ALogP, Molecular Volume, H-bond acceptors, Molecular 3D PolarSASA (see **Supplementary Information**)) able to correlate the loss of affinity of the ATTO Fluo-N-Fab toward tubulin with chemical-physical features of those fluorophores. As reported in **Table 3**, there were clear differences in the size, hydrophilicity profiles, solvent accessibility, and the number of H-bond acceptor atoms between the different fluorescent dyes that might report on their suitability for use in the probes design.

**Table 3.**
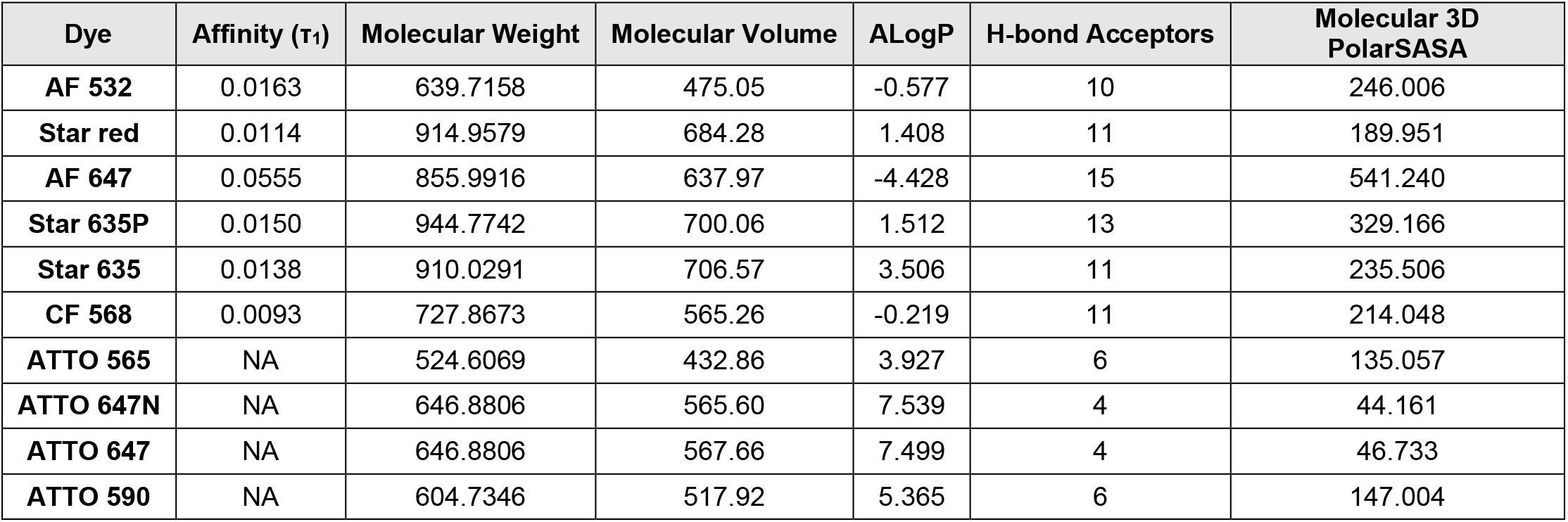

| Dye | Affinity ($\tau_1$) | Molecular Weight | Molecular Volume | ALogP | H-bond Acceptors | Molecular 3D PolarSASA |
| --- | --- | --- | --- | --- | --- | --- |
| AF 532 | 0.0163 | 639.7158 | 475.05 | -0.577 | 10 | 246.006 |
| Star red | 0.0114 | 914.9579 | 684.28 | 1.408 | 11 | 189.951 |
| AF 647 | 0.0555 | 855.9916 | 637.97 | -4.428 | 15 | 541.240 |
| Star 635P | 0.0150 | 944.7742 | 700.06 | 1.512 | 13 | 329.166 |
| Star 635 | 0.0138 | 910.0291 | 706.57 | 3.506 | 11 | 235.506 |
| CF 568 | 0.0093 | 727.8673 | 565.26 | -0.219 | 11 | 214.048 |
| ATTO 565 | NA | 524.6069 | 432.86 | 3.927 | 6 | 135.057 |
| ATTO 647N | NA | 646.8806 | 565.60 | 7.539 | 4 | 44.161 |
| ATTO 647 | NA | 646.8806 | 567.66 | 7.499 | 4 | 46.733 |
| ATTO 590 | NA | 604.7346 | 517.92 | 5.365 | 6 | 147.004 |

To further understand the Fluo-N-Fab affinity towards tubulin, we measured the loss of fluorescence over time in Hela cells stained with the Fluo-N-Fabs able to produce an efficient staining of tubulin by immobilizing cells in a custom-made poly-dimethylsiloxane (PDMS) microfluidic chamber and calculating the fluorescent intensity by confocal microscopy (**see Supplementary Fig 5**). The principle of the assay is to measure the decrease of the intensity of the fluorescent signal with the time under a constant-flow purge of PBS buffer. We evaluated the relative binding strength of each Fluo-N-Fab from the slope of the curve obtained. The relative affinity constants of the six Fluo-N-Fabs were calculated from a bi-exponential fit of the curves (**Figure 2B**), normalized to min= 0 and max=1, following the equation used for fitting the dissociation curves:

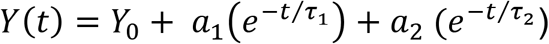

where, *Y(*t*)* is the fluorescent intensity acquired frame by frame, *Y*_0_ is a constant value for the non-specific background (and equals to 0 for normalized curves), T_1_ is the fast decay component associated with the binding strength of each conjugate and T_2_ is the slower decay component associated with the emission of the dye from a non-specific binding intermediate. We used the value T_1_ as a parameter for the measurement of the relative binding strength of the fluorescent conjugate. The assignment of each component is made assuming that the dissociation-dependent component T_1_ is affected only by the flow rate of PBS, while the T_2_ is mainly calculated from the unbound fraction of the Fluo-N-Fab. The calculated T_1_ values for each Fluo-N-Fab are showed in **Table 3**. The binding affinities of the different Fluo-N-Fabs to tubulin were confined to a narrow range, with the Fab_tubulin_ – CF 568 having the highest affinity and the Fab_tubulin_ – AF 647 having lowest affinity amongst the Fluo-N-Fabs tested (**Figure 2B**). However, each of these Fluo-N-Fabs stained endogenous tubulin with no gross differences in staining patterns observed (**Figure 2A**).

Altogether, these experimental data along with the molecular descriptors of the selected fluorophores suggested that a dye should possess a hydrophilicity range ALogP between - 4 and 1, a H-bond acceptor atom number between 10 and 15 and a solvent exposed molecular area ranging from about 200 to 500 A^2^ (**Table 3**) to be suitable for Fab conjugation and to preserve the affinity of the probe for the epitope.

### FAB-FLUOROPHORE CONJUGATES EXHIBIT HIGH PENETRANCE AND ACCESS TO THE EPITOPE IN FIXED CELLS

We utilized these predictive parameters and selected the CF 568 and AF 647 for conjugation with Fab_tubulin_ to determine if this Fluo-N-Fab exhibited enhanced staining properties as compared to conjugating full-length antibodies. To test for this, we analyzed if the Fluo-N-Fabs could penetrate and localize to mid-bodies, these are transient tubulin-rich structures, know to represent highly crowded areas that impairs probe access that form at the sites of abscission between two nascent cells during cell division ^58^ In general, aldehyde-based fixatives crosslink all proteins contained within these tubulin-rich structures, leading to the formation of highly crowded areas that impairs probe access. This is particularly true when either indirect immunofluorescence using secondary antibodies or direct immunofluorescence with primary antibodies techniques are utilized ^59^ ^60^ ^61^. Indeed, the commonly used sandwich format of secondary immunofluorescence prevented the saturation of available sites for epitope recognition in poorly accessible regions of the mid-body (**Figure 2D**). In contrast, the Fluo-N-Fabs generated in this study efficiently stained mid-bodies in cells, revealing an intricate network of filaments within a crowded midbody (**Figure 2D**).

### QUANTITATIVE EVALUATION OF THE LINKAGE ERROR OF FAB PROBES BY STORM IMAGING

Finally, we tested the ability of the Fluo-N-Fabs to reduce the linkage error often observed in SMLM techniques while using primary or secondary antibodies conjugated to fluorophores. Amongst the subcellular structures analyzed to date, the diameter and volume of microtubules, polymers of tubulin within cells, have been precisely measures by electron microscopy that does not require the use of binders and thus can be considered free of linkage error ^62,63^. We hence resorted to analyzing the diameter of microtubules with our Fluo-N-Fabs using *direct* Stochastic Optical Reconstruction Microscopy (*d*STORM). We compared and contrasted the accuracy of spatial resolution and the linkage error generated by the Fab_tubulin_ – AF647 conjugate against the readings obtained while using either a full-length Tubulin antibody coupled to a fluorophore (Ab I) or the full length Tubulin antibody coupled to a fluorophore conjugated secondary antibody. We selected the dye AF647 for these analyses as this fluorophore has greater stability in super resolution microscopy based measurements and does not undergo loss of fluorescence due to rapid blinking. We evaluated the full-width at half maximum (FWHM) of cross sections of microtubules and calculated the median value of microtubule cross sections as a measure of the distance between the fluorescent emission and the biological target due to the probe size (**Figure 3A**, **Table 4 and Supplementary Table 1**).

**Figure 3:**
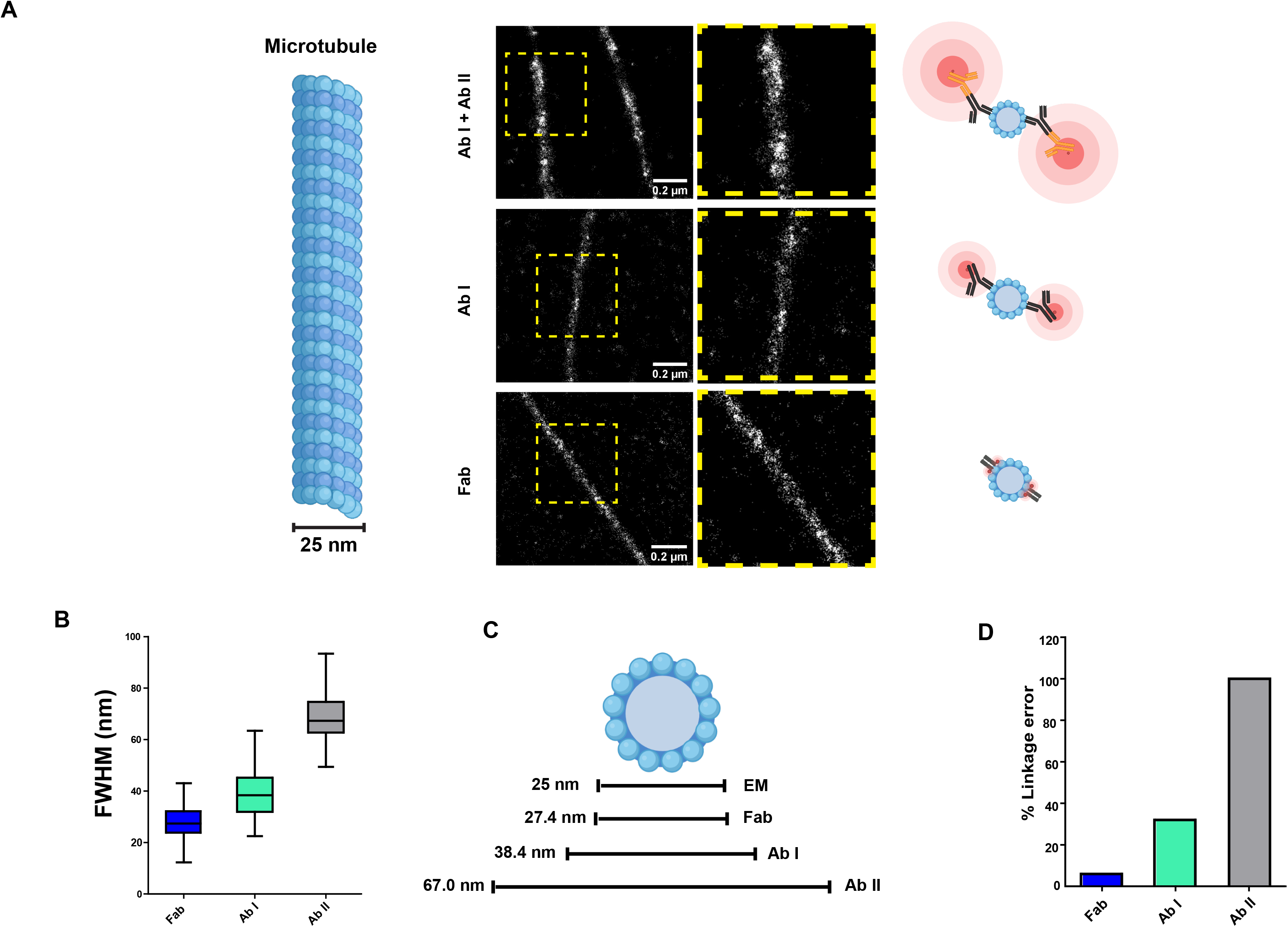
Linkage error produced from Fluo-N-Fabs in STORM microscopy. **A)** Microtubules staining in STORM microscopy obtained using (from up to down) commercially available secondary antibody AF 647, anti-α-tubulin primary antibody randomly labelled with AF 647, anti-α-tubulin AF 647 Fluo-N-Fab; **B)** Fitted values (nm) of the FWHM of microtubule cross sections obtained in 3 independent experiments; **C)** Microtubule external diameter values obtained comparing the electron microscopy value (25 nm) with the one obtained with the N-teminal labelled Fab fragment; **D)** Evaluation of the percentage of linkage error produced from anti-α-tubulin primary antibody randomly labelled with AF 647 and the anti-α-tubulin N-terminal labelled with AF 647.

**Table 4.**

| | Microtubule thickness<br>(fitted values;<br>nm, ± SD) | Spatial displacement (max)<br>(95% CI; nm, ± SD) | Percentage of linkage error<br>(referred to indirect IF) |
| --- | --- | --- | --- |
| Fab | 27.4 | ± 1.19 (2.60) | 6 % |
| Ab I | 38.4 | ± 6.68 (8,36) | 32% |
| Ab II | 67.0 | ± 21.0 (23,0) | - |

The microtubule external diameter measured with the Fab_tubulin_ – AF647 conjugate, the full length primary, and secondary antibodies in these experiments were, respectively, 27.4 ± 1,19 nm, 38.4 ± 6.68 nm and 67.0 ± 21.0 (best fit ±95% confidence interval, average median values over 80 measurements) (**Figure 3B**). These results indicated that the mean external diameter of microtubules measured with the Fab_tubulin_ – AF647 conjugate was extremely similar to the ones measured by electron microscopy, calculated to be 25 nm ^64^ and other STORM measurements with nanobodies ^62^. This significant improvement in nano-structure determination reflects the increased precision by which the molecular coordinates of the fluorophore are localized within 2 nm from the protein target.

We then sought to assess the linkage error produced from the Fab_tubulin_ – AF647 conjugate. We considered that the mean diameter of the microtubule measured by electron microscopy (25 nm) is a value that corresponds to a 0 nm linkage error, our lower limit in the linkage error ruler. As an upper limit, we considered the mean value of microtubule diameter obtained in our STORM experiments using the secondary antibodies that corresponded to

~ 67 nm, in accordance with the measurements reported earlier ^62^. We assumed that these values represented the maximal outliers of the linkage error ruler for microtubules. Based on our STORM measurements, we calculated the relative linkage error obtained from the Fab_tubulin_ – AF647 conjugate. The linkage error produced from N-terminally labelled the Fab_tubulin_ – AF647 conjugate was ~ 2.4 nm (**Figure 3D**). The full-length primary antibody that is randomly labelled had a linkage error of ~13.4 nm and the secondary antibody exhibited an error of ~42nm (**Figure 3D**). When represented as a percentage, the Fab_tubulin_ – AF647 conjugate had a 6% linkage error while the randomly labelled primary antibody produced an error of 32% (**Figure 3C and 3D**).

We note here that the linkage error produced from the Fab_tubulin_ – AF647 corresponds to the 2 nm distance that separates the N-terminal amino group from the antigen binding surface of the Fab. This observation supports the validity of our marking techniques for the production of Fluo-N-Fab. Interestingly, in a previous study ^62^ it was reported that staining the microtubules with anti-α-tubulin nanobodies labelled with AF647 resulted in a FWHM value of 39 nm. This is probably due to the basic pH used for the conjugation of the nanobody. Indeed, the random labelling of the nanobody lysines residue could contribute to increase the off-target of such small binder from the epitope of interest while the site-selective labelling of the N-terminal amino groups in the Fab fragments drastically improve the correct visualization of microtubules as shown by the presented data.

## DISCUSSION

Important limitations common to many super resolution imaging techniques nowadays reside in the properties of the fluorescent probes used for detecting nanostructures, rather than in the optical properties of the microscopes. Thus, an improvement of super resolution imaging analyses hinges upon designing improved probes.

A key challenge is the precision labelling of nanostructures in SMLM by the generation of probes that substantially decrease the linkage error. The linkage error problem is particularly relevant in the case of primary and secondary antibodies that are widely used as epitope binders in super resolution microscopy experiments. They represent the most popular and readily available markers for the detection of cellular structures in biological studies and are chemically conjugated with an uncontrolled number of fluorescent dye molecules. Both classes of markers have inherent problems due to their large size and random distribution of the fluorophores, often generating a spread in the fluorescence signal that usually leads to mislocalization errors and consequentially to the presence of artefacts in super resolution microscopy.

The question of how to overcome the linkage error problems and improve the sensitivity of modern imaging techniques has been waiting for satisfactory answers for some time. Here, we present a simple and reproducible protocol for the production of N-terminal Fab-fluorophore conjugates (Fluo-N-Fabs) with organic dyes that allow for the precision labelling of nanostructures using SMLM. The selective labelling of the N-terminal amino groups of Fab fragments reduces the linkage error by more than five-fold in comparison to conventional reagents, while preserving the numerous and unique properties of antibodies.

The development of a chemical reaction that is able to suitably modify the N-terminal amino groups of proteins is still a complex task. Some of the most popular examples of reactions that have been tried to achieve a specific labelling of the N-terminal amino group of native proteins and antibodies include the site-selective diazotransfer reaction for azide introduction with sulfonyl aziedes ^65^, biomimetic transamination ^66^ and acylation ketene mediated ^67^. However, these reactions require the introduction of specific chemical modification to both the dye molecules and the protein target, constituting an important practical drawback. Additional limitations may be determined by the coupling reaction itself, depending on the specific N-terminal sequence ^68^ and the stability of the conjugated products ^69^ ^70^. Here, we developed a reaction protocol based on the selectively high reactivity of the N-terminal amino groups versus other amino groups at low pH. Thus, selectivity toward the Fab N-terminal amino group is achieved through a fine control of the pH reaction conditions using a simple and mild procedure. We show that when conjugation reactions are performed at a specific pH of 6.5, it is possible to selectively label the N-terminal amino groups of Fab fragments obtained from monoclonal IgG_1_ mouse and polyclonal IgG rabbit antibodies using commercially available N-hydroxysuccinimide esters fluorophores.

Moreover, we defined the criteria for the selection of the fluorophore suitable for the N-terminal labelling of our Fab based probes. We observed that this can be strongly influenced by the hydrophobic properties of fluorophores. Indeed, fluorophores that showed the highest hydrophobic properties (i.e the studied ATTO dyes) irremediably compromise the stability of the Fab fragment and its affinity to the epitope of interest.

We have tested the properties of the Fluo-N-Fabs to reduce the linkage error in SMLM techniques by STORM microscopy. Using the value of the external diameter of tubulin as a molecular ruler, the value of the linkage error observed using our probes was substantially lower to the one found when using conventional labelling reagents such as random fluorescently labelled primary and secondary antibodies. In fact, we estimate that the linkage error of the Fluo-N-Fabs is by far the lowest that has been reported so far and is strikingly close to the 2 nm distance that separates the labelled N-terminal amino group from the antigen binding domain surface of the Fab.

Notably, the reduction of the linkage error in SMLM can be achieved not only by reducing the size of the linker but also by the introduction of a controlled number of fluorophores (one in our case) in fluorescent probe. A similar concept has been recently proposed in the interesting work of Früh et al. ^71^ where these authors propose the introduction of a controlled number of fluorophores on the hinge region of the F_c_ part of full-length antibodies. These types of probes also reduce the linkage error of the primary full-length antibodies, but the calculated distance between the F_c_ fragment of the antibodies and the antigen of interest reported in their studies is still larger than in our experiments using the Fluo-N-Fabs. Thus, conjugating the N-terminus of a Fab that is closest to the epitope binding zone results in a ‘zero-distance’ label that is quite possibly the lowest linkage error that can be achieved without interfering with the binding affinity of probe.

Additionally, another advantage of our Fluo-N-Fab system is that the Fab fragment is sufficiently small, in striking contrast with full length antibodies, to significantly improve tissue penetration and access to epitopes of interest in crowded cell compartments.

As already noted, our choice to use Fab fragments for the generation of our probes is linked to the numerous advantages of these types of reagents. The most important of them is the very wide and practically complete range of commercially available antibodies from which it is possible to generate Fab fragments matching the specific research project. In sum, these myriad features of the Fluo-N-Fab reagents can be further exploited in other potential applications like the staining of endogenous proteins expressed in cells or tissues isolated from patients.

We believe that in addition to the minimization of the linkage error, the use of Fluo-N-Fabs should significantly broaden the experimental design of the many laboratories that are focusing on the application of advanced fluorescent imaging to assess important biological questions.

The minimal linkage error of Fluo-N-Fabs will certainly boost the capabilities of modern super-resolution microscope technologies (SMLM and MINIFLUX^3^), which already today can localize a fluorophore with an error well below 10 nm, by enabling the most precise localization of the target available today. Furthermore, the small size of the Fluo-N-Fabs enables an improved sample penetration and thereby the detection of the targets in densely packed cellular environments. This widens the range of Fluo-N-Fabs applications to all fluorescent imaging techniques, which require small sized and versatile affinity reagent.

## METHODS

### N-TERMINAL LABELLING OF MONOCLONAL ANTIBODY AND FAB FRAGMENTS

The Fab were obtained by enzymatic digestion with Ficin (Immobilized Ficin, Thermofisher) or Papain (Immobilized Papain, Thermofisher) of the full length antibodies following the manufacturer instructions. The purified Fabs were is buffer exchanged against 10 mM phosphate solution buffered at pH of 6.50 and concentrated using the Speed Vacuum (SAVANT Speed Vacuum, Thermofisher) until the final concentration of 5 mg/mL. For the labelling reaction we used N-hydroxysuccinimide dyes that we found be suitable for our purposes. The dye solution is prepared dissolving the dye powder in anhydrous N,N-dimethylformamide (DMF, Sigma-Aldrich). The dye to protein molar ratio was adjusted between 2 - 5 in order to provide a DOL around 1, and the molar concentration of reactive NHS esters in DMF was such that no more than 5% of the entire reaction volume consisted of organic solvent. The dye solution in DMF is added to the protein solution and the reaction is kept in a water bath at 37°C for 1h. After, to each reaction mixture was added the quenching solution containing 2 M glycine, 1% Triton-x 100 or Tween-20 at the pH of 8-8.6. The amount of quenching solution is at least 3 volumes of reaction solution. The quenching step is maintained at 37°C for 15 minutes and then shifted at 4°C overnight. It is very important avoid any strong light capable of photo-bleaching the fluorophore and for all the steps gentle vortexing of the solution is randomly done.

### DEGREE OF LABELLING DETERMINATION

The degree of labelling (DOL) is the average number of dye molecules coupled to a protein molecule. The DOL can be determined from the absorption spectrum of the labelled protein. For the DOL determination, the protein conjugate solution should be completely free from the free dye contamination. Then, the UV/VIS spectrum of the solution is acquired in a window starting at ~250 nm and covering the complete absorbance band of the coupled dye (for our samples at least until 700 nm). The absorption spectra must be acquired using a UV transparent quartz cuvette. The following values has been collected from the absorption spectrum:λ_max_: the wavelength where the absorbance has its maximum (should be close to the absorbance maximum of the pure dye, compare the respective data sheet), A_max_: the absorbance at λ_max_, A_280_: the absorbance at 280 nm. The DOL is calculated from the measured values according to the following equation:

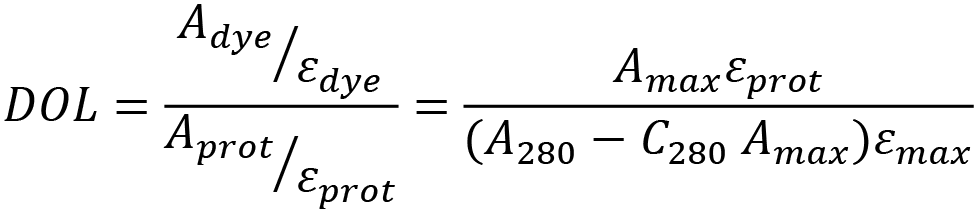

Here, the symbols denote: A_prot_: absorbance of the pure protein at 280 nm (absorption maximum of proteins), ε_prot_: molar extinction coefficient (in M^−1^cm^−1^) of the pure protein at 280 nm,ε_max_: molar extinction coefficient (in M^−1^cm^−1^) of the dye at its absorbance maximum, C_280_: correction factor which depends on the dye’s spectral properties. (the major part of the dyes are showing some absorbance also at 280 nm). The ε_prot_ considered for the determination of the DOL of Fab and full length antibodies are 70000 M^−1^ cm^−1^ and 210000 M^−1^ cm^−1^ respectively. The λ_max_, ε_dye_ and C_280_ for all the fluorophores used in this studies are so listed: Alexa Fluor 647 (Invitrogen, Cat# A20006): 650 nm, 239000 M^−1^ cm^−1^, 0.03; Alexa Fluor 532 (Invitrogen, Cat# A20001): 532 nm, 81000 M^−1^ cm^−1^, 0.09; CF 568 (Biotium, Cat# SCJ4600027): 562 nm, 100000 M^−1^ cm^−1^, 0.08; Star 635p (Abberior, Cat# ST635P): 638 nm, 120000 M^−1^ cm^−1^, 0.4; Star 635 (Abberior, Cat# ST635): 635 nm, 110000 M^−1^ cm^−1^, 0.38; Star Red (Abberior, Cat# STRED): 638 nm, 120000 M^−1^ cm^−1^, 0.32; ATTO 565 (ATTO-TEC, Cat# 72464): 564 nm, 120000 M^−1^ cm^−1^, 0.12; ATTO 590 (ATTO-TEC, Cat# 79636): 593 nm, 120000 M^−1^ cm^−1^, 0.43; ATTO 647 (ATTO-TEC, Cat# 18373): 647 nm, 120000 M^−1^ cm^−1^, 0.04; ATTO 647N (ATTO-TEC, Cat# 18373): 646 NM, 150000 M^−1^ cm^−1^, 0.03.

### MASS SPECTROMETRY SAMPLE PREPARATION: *in situ* DIGESTION OF ANTI-C-MYC FAB

The reaction products of anti-c-Myc Fab labelled with the CF 568 dye were purified using Size exclusion chromatography (SEC-1 MAbPac, Thermofisher) (**Supplementary Fig. 6A**) followed by Hydrophobic interaction chromatography purification (HIC-10 MAbPac, Thermofisher) (**Supplementary Fig. 6B**). The anti-c-Myc labelled Fab was collected and used to check the conjugation reaction selectivity toward the N-terminal by mass spectrometry. The Fab was first reduced with DTT and alkylated with Iodacetamide at room temperature in the dark. Subsequently the two chains were separated by SDS-PAGE on 15% polyacrylamide gel. Comassie stained protein bands corresponding of the light and heavy chains were excised from the gel and washed in 50mM ammonium bicarbonate pH 8.0 in 50% acetonitrile to a complete distaining. Gel pieces were re-suspended in 50mM ammonium bicarbonate, pH 8.0, reduced with 10mM DTT and alkylated with a 55mM solution of iodoacetamide at room temperature in the dark. The excess of reagent was discarded, the gel pieces were washed several times with the same buffer, resuspended in 50mM ammonium bicarbonate and incubated with trypsin. The supernatant containing the resulting peptide mixtures was removed and the gel pieces were re-extracted with acetonitrile in order to remove all the peptides still present in the gel. The two fractions were then collected and freeze-dried.

### MASS SPECTROMETRY ANALYSIS OF ANTI-C-MYC FAB: MALDI-MS ANALYSIS

MALDI-MS analyses were carried out on a 4800 plus MALDI TOF-TOF mass spectrometer (AB Sciex) equipped with a reflectron analyser and used in delayed extraction mode with 4000 Series Explorer v3.5 software.

For the analyses, 0.5μl of peptide mixture were mixed with an equal volume of a-cyano-4-hydroxycynnamic acid as matrix (10 mg/ml in 0.2% TFA in 70% acetonitrile), loaded onto the metallic sample plate and air dried. Mass calibration was performed using the standard mixture provided by manufacturer. MALDI-MS data were acquired over a mass range of 600–5000 m/z in the positive-ion reflector mode. MS spectra were acquired and elaborated using the software provided by the manufacturer.

### MASS SPECTROMETRY ANALYSIS OF ANTI-C-MYC FAB: LC-MS ANALYSIS

The triptic peptides were analyzed by a LTQ XL-Orbitrap ETD equipped with a HPLC NanoEasy-PROXEON (Thermo Fisher Scientific). They were loaded, concentrated and desalted on a C18 Easy-Column (L = 2 cm, ID = 100 μm; cat. no. 03-052-619, Thermo

Scientific SC001). They were then fractioned on a C18 reverse-phase capillary column (L = 20 cm, ID = 7.5 μm; cat. no. NS-AC-12, Nano Separation, Niewkoop, Netherlands) at a flow rate of 250 nl/min in a gradient from 5 % to 95 % buffer B (eluent B: 0.2 % formic acid in 95 % acetonitrile; eluent A: 0.2 % formic acid and 2 % acetonitrile in ultrapure water) over 87 min.

The mass spectrometric analyses provided MS and MS/MS spectra of the peptides. Mass spectra data were then used to search a non-redundant protein database by using the Mascot software.

The files generated by the analysis processing were analyzed by Mascot. The light chain was identified with a score of 1548 while the heavy chain with a score of 1156. The controlled sequence portions are highlighted in The results of the mass analyzes led to the verification for the light chain of a sequence coverage equal to 100% while for the heavy chain a sequence coverage equal to 64.5%.

### CELL CULTURE

Immuno-fluorescence procedures and dissociation curves were performed by confocal microscopy in HeLa cells. Measurements of α-tubulin mean diameter were performed by Single Molecule Localization Microscopy (STORM) in U-2 OS cells.

HeLa cells were purchased from American Tissue Type Collection (ATTC, USA), and were grown in Dulbecco’s modified Eagle’s Medium (DMEM), supplemented with fetal bovine serum (FBS), penicillin, streptomycin and L-glutamine (all from Gibco/BRL, NY, USA). Trypsin- EDTA was used to detach and split the cells. All microfluidic experiments are done with HeLa cells.

U-2 OS cells were purchased from American Tissue Type Collection (ATTC, USA), grown and maintained in phenol-red free DMEM (Invitrogen) supplemented with 10% FCS (Labforce), Glutamax (Invitrogen), without any antibiotics added. Cells were plated on 18 mm round coverglass (Cat# 1.5H, 170 μm ± 5 μm precision cover glasses, Fisher Scientific, Thermo Fisher) and after 24 h were gently fixed with a EM Grade 4% paraformaldehyde aqueous solution (from 8% sealed ampules, EM Science) dissolved in cytoskeleton stabilizing buffer (CSB), previously developed by Whelan et al. to prevent image artefacts, consisting of 500mM NaCl, 80 mM PIPES, 1 mM MgCl_2_, 1 mM EGTA, 2mM sucrose, pH 6.2 ^72^. Cells were then permeabilized for 30 minutes at room temperature in a 0.05% w/v saponin solution dissolved in PBS (Sigma, quillaja bark powder), and then washed 3X with PBS before being incubated with the staining reagents.

Mycl-9E10 hybridoma cells were purchased European Collection of Authenticated Cell Cultures (ECACC). Cells were cultured in RPMI 1640 with 2 mM Glutamine and 10% Fetal Bovine Serum Low IgG (Invitrogen), we used this particular kind of serum in order to decrease the contamination of anti-c-Myc antibody from any other antibody sources. Cells were cultured between 3-9×100000 cells/ml; 5% CO2; 37°C. When recovering hybridoma cultures from frozen it is not unusual for growth to be slower than expected initially and there may be an observed decrease in viability. Establishment of an actively proliferating culture may take up to 2 weeks. Once the culture is established in culture the FBS can be reduced gradually until 2%. The supernatant has been collected and used for the production of the anti-c-Myc antibody (IgG_1_ mouse monoclonal, κ isotype).

### EVALUATION OF THE IMPACT OF FLUOROPHORES ON THE BINDING STRENGHT OF FAB FRAGMENTS

Six of the ten fluorescent Fab conjugates prepared in this study were suitable for staining the cytoskeleton of HeLa cells. For the same epitope-binder combination, we evaluated the impact of each fluorophore on the binding strength of the conjugate by measuring the dissociation rate during a constant wash out. HeLa cells were grown on custom-made PMDS microfluidic chamber (described in **Supplementary Fig 6**), and are fixed in 4% paraformaldehyde PBS solution buffered at pH 7.00, washed three times with PBS and permeabilized with a PBS solution of saponin at 0.05% w/v (later named washing buffer). The same surfactant concentration is maintained during the entire experiment to keep the cells continuously permeabilized in a purge flow of washing buffer. The chamber is constantly held at a fixed position from the objective to avoid fluctuations around the on-focus position, as movements of the sample from the initial focal plane would change the emission intensity; drifts along Z are minimized by the built-in autofocus module of the microscope (Adaptive Autofocus, Leica SP5-II). For each fluorescent Fab conjugate, the bleaching rate is evaluated before starting the acquisition to check for possible artefacts due to serial acquisitions, and therefore excitation laser power is adjusted to avoid random decreases in the emission intensity independently from the wash-out, which are not related to the dissociation of the antigen-Fab couple but rather to the hydrophobicity of the dye ^73^. Frames are acquired every 5 seconds, during a time lapse of 200 seconds, and the fluorescence emission intensity is recorded within a ROI including the cell cytoskeleton. The background intensity of unbound staining is excluded from the analysis, because its intensity is not dependent on the dissociation equilibrium.

### SAMPLE PREPARATION FOR SINGLE MOLECULE LOCALIZATION MICROSCOPY

Covers were rinsed from PBS and incubated for 60 min with 5 μg/ml AF647-labeled Fabs / primary antibody in blocking buffer (0.05% saponin in PBS). The antibody used in this study is a mouse monoclonal antibody directed against α-Tubulin (clone DM1A, Sigma), labelled or cleaved into its Fabs as described in the section above. For super-resolution SMLM-imaging using the dSTORM / GSDIM protocol, 18 mm coverslips (50000 cells/slide) were stored in PBS after fixation and immunolabelling at 4°C. The coverslips were mounted onto a single depression slide (76 mm × 26 mm) and the cavity filled with freshly prepared 90-100μl GLOX-MEA buffer (0.5 mg/ml glucose oxidase (Sigma-Aldrich, Cat# G7141, 40 μg/ml catalase (Sigma-Aldrich) 10% w/v glucose (Sigma-Aldrich) 50 mM Tris-HCl pH 8.0, 10 mM NaCl and 10 mM β-mercaptoethylamine (Sigma-Aldrich, Cat# M9768-5G)). MEA solutions were freshly dissolved before each use, and the ideal concentration was determined to be 20mM ±10mM by assessing the blinking frequency during the acquisition of SMLM images. The pH of the highly basic MEA stock solution is adjusted dropwise with conc. HCl using colorimetric indicators due to the electroactivity of MEA towards glass electrodes of common pH-meters. The coverslip is sealed to the depression slide with two-component silicone-glue Twinsil® (Picodent, #13001000) to avoid atmospheric oxygen quenching the fluorophore’s triplet state, which results in a decrease of the fluorophore blinking kinetic.

### SMLM imaging with dSTORM/GSDIM

Imaging was performed with a Leica SR GSD system using a HC PL APO 160× / NA 1.43 oil objective. The images were recorded with an Andor iXon 897 EMCCD camera at 40 Hz using a central 180 pixel x 180 pixel subregion. For excitation, a 532 nm laser (500 mW maximum power output) and a 642 nm laser (500 mW maximum power output) were used and attenuated using an AOTF when appropriate. The two fluorophores were recorded sequentially and image acquisition, single molecule analysis and image reconstruction was performed with Leica LAS X 1.9.0.13747.

