## Supplementary for "Precision localization of cellular proteins with fluorescent Fab-based probes"

*Equal contribution

### Supplementary materials

#### Supplementary materials #1: *In silico* prediction of amino-group pKa values

The pKa values of the amino groups (both N-terminal residues and the lysines) were predicted using two computational applications, namely Qsite [1 2] and Potein Titration Curve, contained respectively in the Small-Molecule Drug program suites Discovery Suite and Biologics Suite, distributed by Schrodinger (Small-Molecule Drug Discovery Suite 2018-2, Schrödinger).

Before that, the following preparatory steps were carried out for each investigated FAB: i) download of the experimentally solved structure from the PDB database (www.rcsb.org), selecting the most complete structure, consistent with the respective primary sequence; ii) optimization of the structure by using the Protein Preparation Wizard tool (Schrödinger Release 2018-2: Schrödinger Suite 2018-2 Protein Preparation Wizard), here, any proteins/molecules co-present in the crystal was removed, as well as the solvent and any ions and metals. Then, the hydrogen atoms were added, any ambiguous orientations of the side chains were resolved and the protonation state generated at a pH of 7.4.

The resulting refined structures of the FABs are processed with Qsite, a hybrid of quantum mechanics (QM) and molecular mechanics (MM) software, with the intent of elaborating more finely the geometry and energies of the N-terminal residues and of all lysines present, predisposing in this to a more precise calculation of the pKa values. The calculation is carried out as follows:

1) a dedicated calculation is carried out for each amino group.

2) The terminal residues are treated entirely in the QM way, while for the amines of the lysines only the atoms of the side chain are treated. The remaining atoms of the FAB under examination are subject to MM calculation.

3) For each amino group, a spherical neighbourhood of 12 Armstrong radius, centred on the group under examination, defines the atoms for which non-bonding interactions are taken into account.

4) Calculation details:

• The calculation strategy applied is the DFT-B3LYP [3 4]

• The force field for the MM phase used is OPLS_2005 [5]

• The applied dielectric constant is equal to 80 (thus simulating the aqueous environment)

• Molecular constraints are imposed on the backbone of the FAB, so as to limit any unfolding phenomena during the energy minimization cycles of the System.

• The energy convergence values for minimization are i) the energy value and ii) the energy gradient, set to 0.1 kcal / mol and 0.01 kcal / (mol Å) respectively; minimization algorithm is the "Truncated Newton".

The optimization performed with Qsite is followed by the prediction of the pKa values of the amic group being analyzed with Potein Tirtration Curve, which uses the PropPKA algorithm [6 7]. The values thus calculated separately are collected and reported for each analyzed Fab.

**Supplementary materials #2: Light chain N-terminal amino group of Fab 2OR9 protonation at pH 6.5**

The in silico calculation of the amino groups pKa values in the Fab 2OR9 showed that the pKa value of the light chain N-terminal amino group was significantly higher than expected (pKa= 9.96) while the one of the heavy chain was 7.8. This peculiarity may find explanation by looking at the crystal structure of the Fab itself, where the N-terminal amino group may interact with two glutamic side chains at positions 27 and 93 of the light chain (**Figure S1A**), stabilizing the protonated form by salt bridges. This coordination might result in an unusually high pKa value for that N-terminal amine, which becomes unavailable for the labelling reaction at a neutral pH. We analysed 124 amino acid sequences of variable domain $IgG_1$ mouse k isotype antibodies sequences available in the IMGT database, in order to understand if the coordination of the N-terminal amino group with glutamic acid is frequently occurring. We generated a WebLogo of the aligned sequences (**Figure S1B**) and we found that while the glutamic acid 27 was usually highly conserved instead of the glutamic acid 93 was always absent. We also considered the folding of the $IgG_1$ monoclonal mouse Fabs analysed, in particular 16 Fabs crystal structures have been selected according to the criterion of maximizing the diversity of the amino-acid sequence and to exclude a possible amino acidic-structural bias. We calculated the pKa values of the N-terminal amino groups of the heavy and light chain of those Fabs crystal and we observed that only few of them were showing values similar to the one of the light chain Fab 2OR9 (**Table 3**). This analysis shows that in most of the Fab considered, the respective pKa values of the two chains show a mean value of 7.5. Nevertheless, in a smaller number of cases one of the two NH2 terminal groups can show a significantly higher pKa value. In this particular situation there is only one N-terminal amino group actually available for the chemical modification and, as for the selectivity of the reaction, this is related to the particular pH that is chosen for the reaction. Indeed, the goal of these pKa statistical analyses was to understand how to select a pH value for the labelling reaction that will take in consideration the relationship between selectivity of the reaction and availability of the amino groups.

**Supplementary materials #3: Optimization of the Fab-dye conjugation protocol**

Fab were obtained by enzymatic digestion of commercially available monoclonal mouse antibodies and polyclonal rabbit antibodies, namely anti-α-tubulin, anti-Pi4K(III)β, anti-ARF, anti-BARS and anti-14.3.3γ. After specific purification steps, Fabs were then reacted with AF 647 N-hydroxysuccinimide ester at pH at 6.5, which was considered particularly appropriate for a selective modification of the N-terminal. After 1 hour at 37 °C, the reaction mixtures were treated with an excess of ethanolamine to quench the excess of amino reactive dye. All the labelled Fab were purified by means of gel filtration chromatography and stabilized by the addiction of BSA. The labelled products were finally tested by immunofluorescence. Interestingly, the monoclonal (mouse) and polyclonal (rabbit) Fabs, behave differently. In particular, at variance with polyclonal Fabs that were able to produce an optimal staining of their cellular targets, the monoclonal Fabs exhibited unexpected non-specific signals.

In the attempt to understand the cause of this difference, we have compared the mouse monoclonal Fabs with rabbit polyclonal Fabs labelled with AF 647 by SDS-PAGE gel (20%)

(**Figure S2A**). In both cases, The SDS-PAGE revealed the presence of various fluorescent polypeptides and proteins, including some BSA (in some lanes). Although BSA was added as stabilizing agent to the Fab after the labelling reaction, it is very likely that its labelling was due to a residual N-hydroxysuccinimide ester) entrapped into purified Fabs and still capable of react. If so, this was an indication that the quenching step with ethanolamine [8] (after the reaction was inadequate. It is interesting to note that the monoclonal, in respect to the polyclonal Fabs retained a larger amount of dye. To get a better insight into this, we performed three parallel labelling reactions using two different mouse monoclonal Fabs targeting the proteins PI4K(III)β and ARF1, respectively. We changed also the quenching and storage conditions. Glycine containing a surfactant replaced ethanolamine. According to observations by others, the surfactant may possibly act as a phase transfer catalyst. Its presence may provide better ester scavenging activity for hydrophobic dyes in concentrated protein solutions that may have a colloidal property. The fluorescent Fabs were analysed by SDS-PAGE. We compared the results with the homologous monoclonal mouse Fab obtained with previously established quenching conditions (ethanolamine, no surfactant). The bands were analysed using a Typhoon scanner (**Figure S2B**). Overall, these experiments indicated that both ethanolamine and even glycine, if used in absence of surfactant were poor scavengers as labelled BSA was invariably detected. At variance, the addition of glycine in the presence Tween 20 or Triton X-100 (as shown in lanes 3 and 4 of **Figure S2B**) granted the best conditions for quenching (as BSA does not appear to be labelled) and even for storage.

The staining performances were finally tested by confocal microscopy. The results obtained by using glycine/surfactant as scavenger, were superior as the staining of tubulin and PI4KIIIβ was fully void of non-specific signals (**Figure S2C**).

### Supplementary materials #4: *In silico* calculation of fluorophore physic-chemical parameters

Each fluorophore molecule was computed in Schrodinger Maestro and standardized according to the following procedure: 1) generation of the 3-D structure, 2) attribution of the OPLS3 force field [9], 3) selection of the favoured ionization form at pH 7.4, 4) generation of the possible conformational isomers with the MCMM [10] method and 5) selection of the most energetically favoured conformer. A carboxy-metil group was inserted instead of the dioxopyrrolidine group, to reduce the most the unwanted contribute of atoms that actually exits during the functionalization reaction.

A wide selection of molecular descriptors was calculated by means of utilities included in the Biovia Pipeline Pilot software. For each compound the staining effectiveness (positive or negative) and its affinity, as measured by confocal microscopy, are reported. A two-tailed, heteroskedastic Student t test is performed for each descriptor, in order to evaluate in terms of statistical significance (p-value) the hypothesis for which the average values of the two populations differ significantly. p-value threshold for positivity was set at 0.05 (Supplementary **Table 2** sheet 1 available for download in **Supplementary data**).

Furthermore, in order to make the selection of discriminating variables more restrictive, also taking note of the narrowness of the number of samples in the two populations, only the descriptors in which the ranges of values, calculated as the mean value +- one standard deviation, do not overlap are selected (**Supplementary Table 2**, sheet 2).

Subsequently, the experimental affinity values, as measured between the labelled antibody and the antigen were evaluated in terms of linear correlation (r) to each descriptor. Considering the small number of samples in the class under examination, we impose the selection of only the descriptors that have an absolute value correlation> 0.8. Of these, the coefficient of determination (r2) is then calculated as a measure of the between the variability of the data and the correctness of the statistical model used, and relative statistical significance (p-value) (**Supplementary Table 2**, sheet 3). As before, a p-value threshold value of 0.05 is imposed, above which the correlation is deemed not significant. **Supplementary Table 2** sheet 4 resumes the selected molecular descriptors.

**Supplementary materials #5: Evaluation of the impact of fluorophores on the binding properties of N-terminal labelled Fab fragments**

Measurements have been performed in a gas-permeant microfluidic chamber realized in PDMS by soft-lithography (**Figure S6A and S6B**). Then, it was bonded to a common coverslip for confocal microscopy (thickness 150 μm ± 20 μm) by the standard procedure based on O2 plasma. All the measurement of the Fab probes binding has been done under continuous flow operation using a programmable syringe pump (Scientific™ KDS 200P) (**Figure S6B**). Antigen targets were endogenously expressed in HeLa cells. The assay was performed in such a way that solution-phase antibodies, in a dynamic equilibrium with those in binding with the epitopes, were constantly washed-out by a purge flow of PBS buffer (**Figure S6**). Intracellular fluorescence densities for the same Fab-antigen complex (Fab from DM1A antibody and α-tubulin of HeLa cells) are recorded in real time during the dissociation, by scanning confocal mode. The system precisely measures the fluorescence emission of each Fab-dye conjugate excited by an illuminating optical beam, and its decrease proportional to the dissociation of each Fab-antigen conjugated with the six fluorophores which were found to be functional. The fluorescence emission data, acquired from specific ROIs within the cytoskeleton, yield binding curves that are subsequently used to extract a *relative* binding kinetic index of each dye-Fab conjugate, which reflects the specific avidity resulting from dye that is attached.

### Supplementary figures

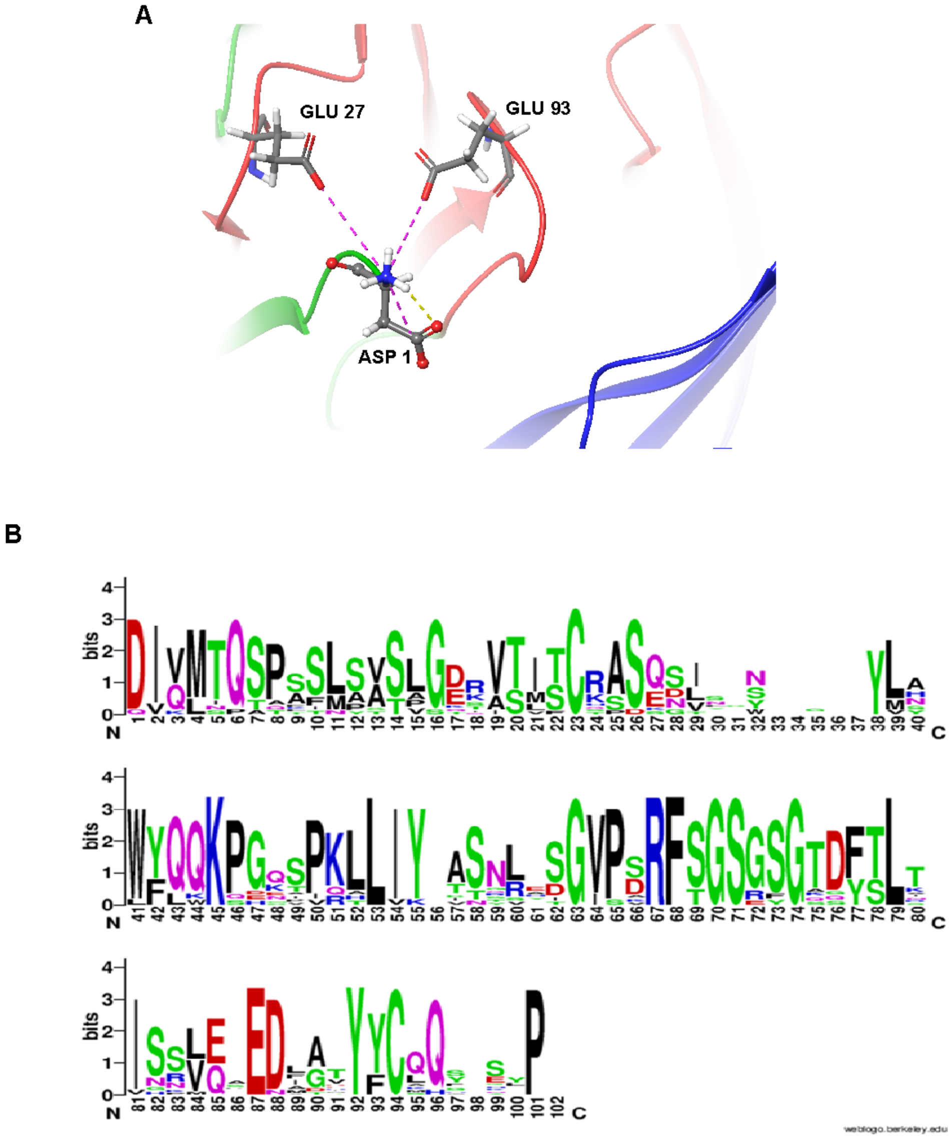

**Supplementary Fig 1: Structural analysis of the Fab 2OR9 light chain and frequency analysis of conserved residues in mouse IgG1 antibodies:** A) Light chain N-terminal amino group of the 2OR9 Fab interacting with the Glutamic acid 27 and 93 through salt bridges interactions (pink) and hydrogen bond (yellow) B) Weblogo light chain Vk domains of IgG1 mouse monoclonal Fab fragments listed in IMGT data base. The logo logo consists of stacks of amino acids symbols, one stack for each position in the sequence. The overall height of each stack indicates the sequence conservation at that position, while the height of symbols within the stack indicates the relative frequency of each amino or nucleic acid at that position.

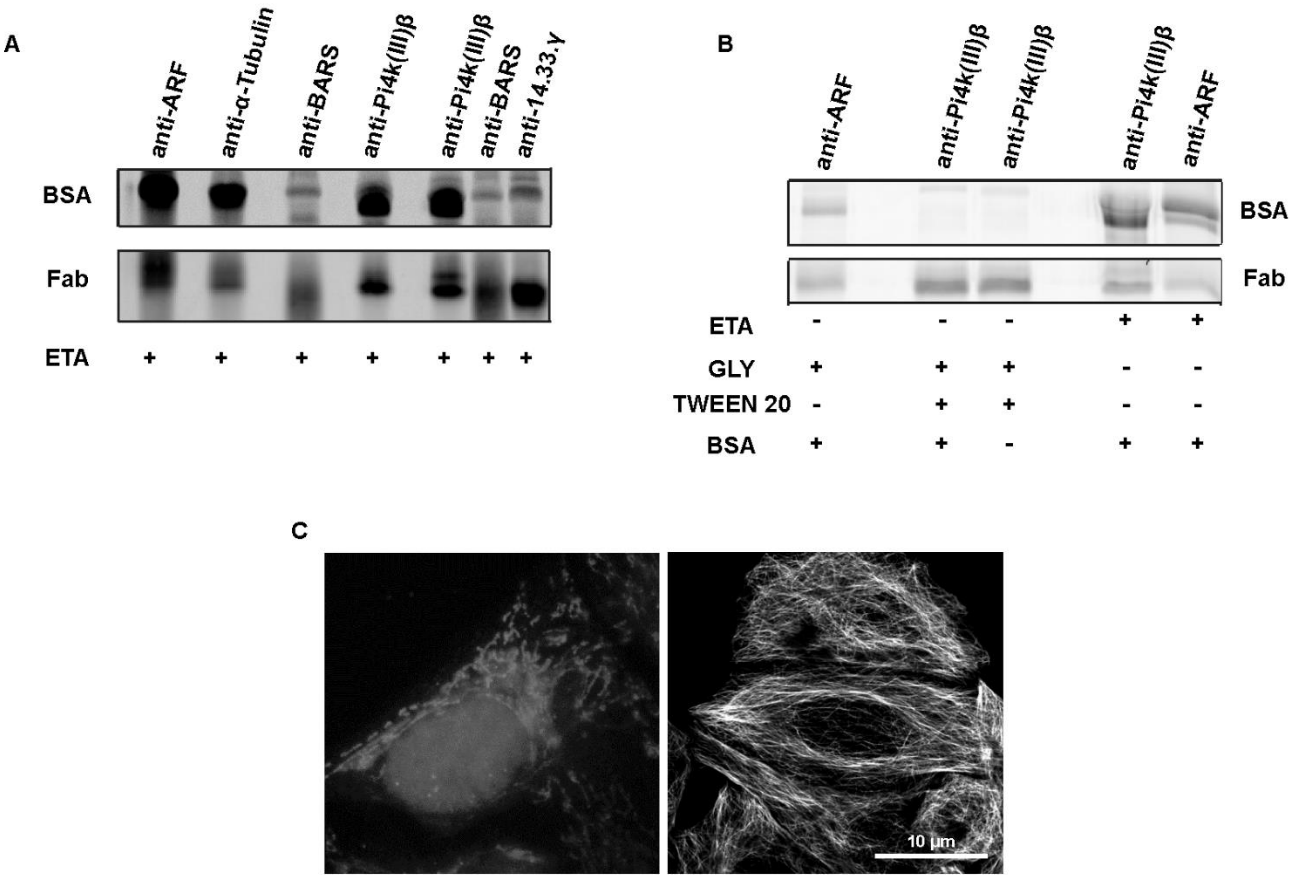

**Supplementary Fig 2: Fab fragments N-terminal labelling protocol**. **A)** Fluorescence emission BSA (66 kDa) and Fabs (HC≈25 kDa, LC≈24 kDa) labelled with AF 647 separated by SDS page. The mouse monoclonal Fab fragments are anti-ARF, anti-Pi4k(III)β and anti-α-tubulin. The polyclonal Fab fragments are anti-BARS and anti-14.3.3γ. All of the Fab fragments are obtained using ethanolamine. The image has been acquired with Typhoon scanner (Amersham) **B)** Comparison between the ethanolamine and the glycine/surfactants quenching protocol. Fluorescence emission of the BSA and Fabs labelled with AF 647 has been acquired with Typhoon scanner (Amersham). **C)** Tubulin staining of Hela M cells (Scale bar 10 μm), on the left is shown the non-specific signals obtained using the scavenger system based on ethanolamine on mouse monoclonal Fab like anti-α-tubulin; on the right is shown the tubulin staining using the scavenger system based on glycine/surfactants.

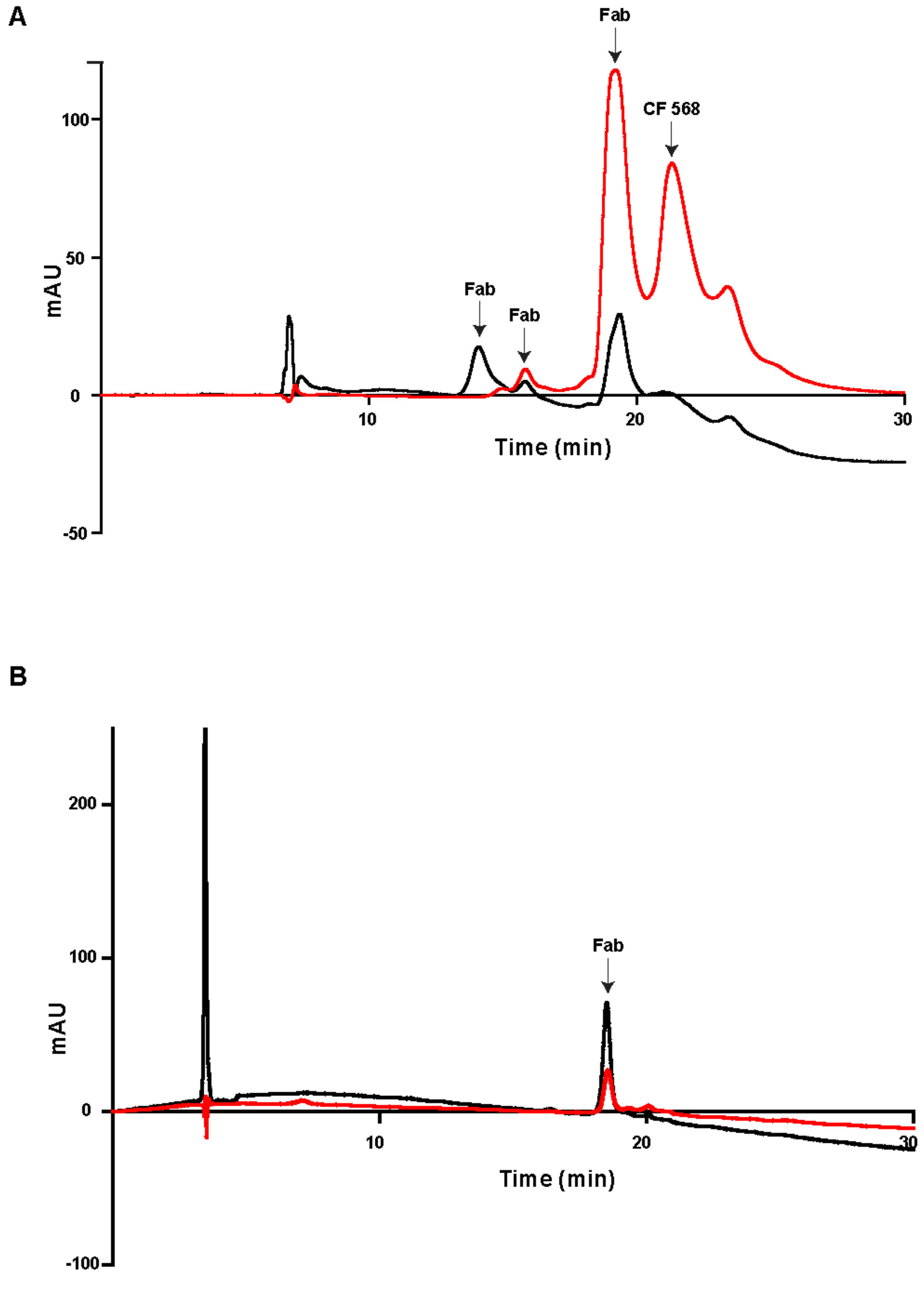

**Supplementary Fig 3: Hydrophobic interaction chromatography analysis of the anti-α-Tubulin Fab and anti-c-Myc Fab labelled with CF 568. A**) HIC-20 chromatogram of anti-α-Tubulin Fab labelled with CF 568 dye. The detection was obtained considering the 280 nm (black) and 560 nm (red) wavelengths for the absorbance of the protein and the fluorophore respectively**. B)** HIC-10 chromatogram of labelled anti-c-Myc Fab labelled with CF 568 dye. The protein and the fluorophore absorbance were detected at 280 nm (black)

and 560 nm (red), respectively.

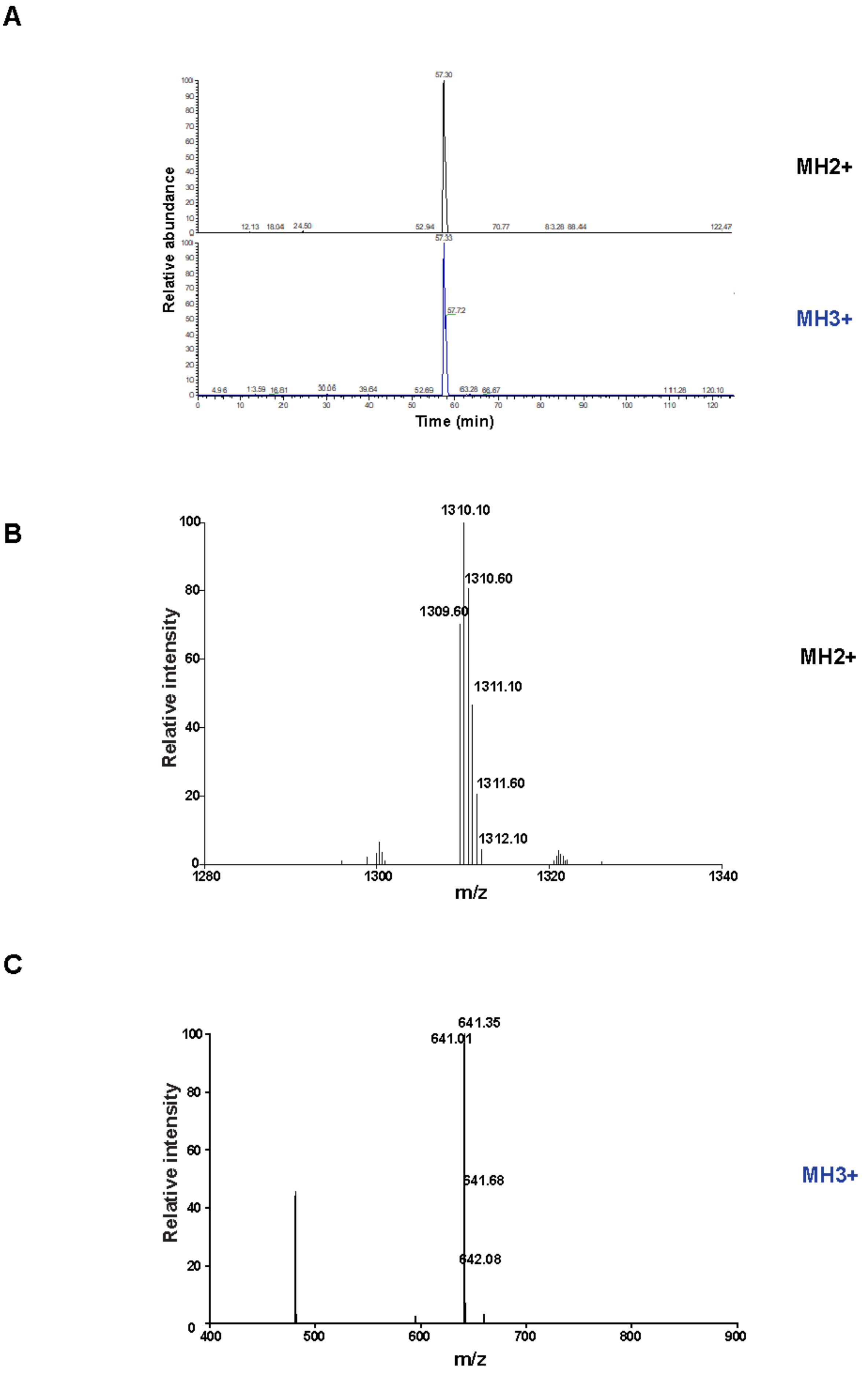
A
Relative abundance
57.30
MH2+
57.33
57.72
MH3+
Time (min)
B
Relative intensity
1310.10
1310.60
1309.60
1311.10
1311.60
1312.10
MH2+
m/z
C
Relative intensity
641.35
641.01
641.68
642.08
MH3+
m/z

**Supplementary Fig 4: Mass spectra of anti-c-Myc Fab heavy chain N-terminal peptide labelled with CF 568 dye. A)** LC-MS Total Ion Chromatogram (TIC) of the HC N-terminal labelled peptide **B)** Mass spectra of the labelled peptide with double charge (MH2+) detected by LC-MS**. C)** Mass spectra of the labelled peptide with triple charge (MH3+) detected by LC-MS.

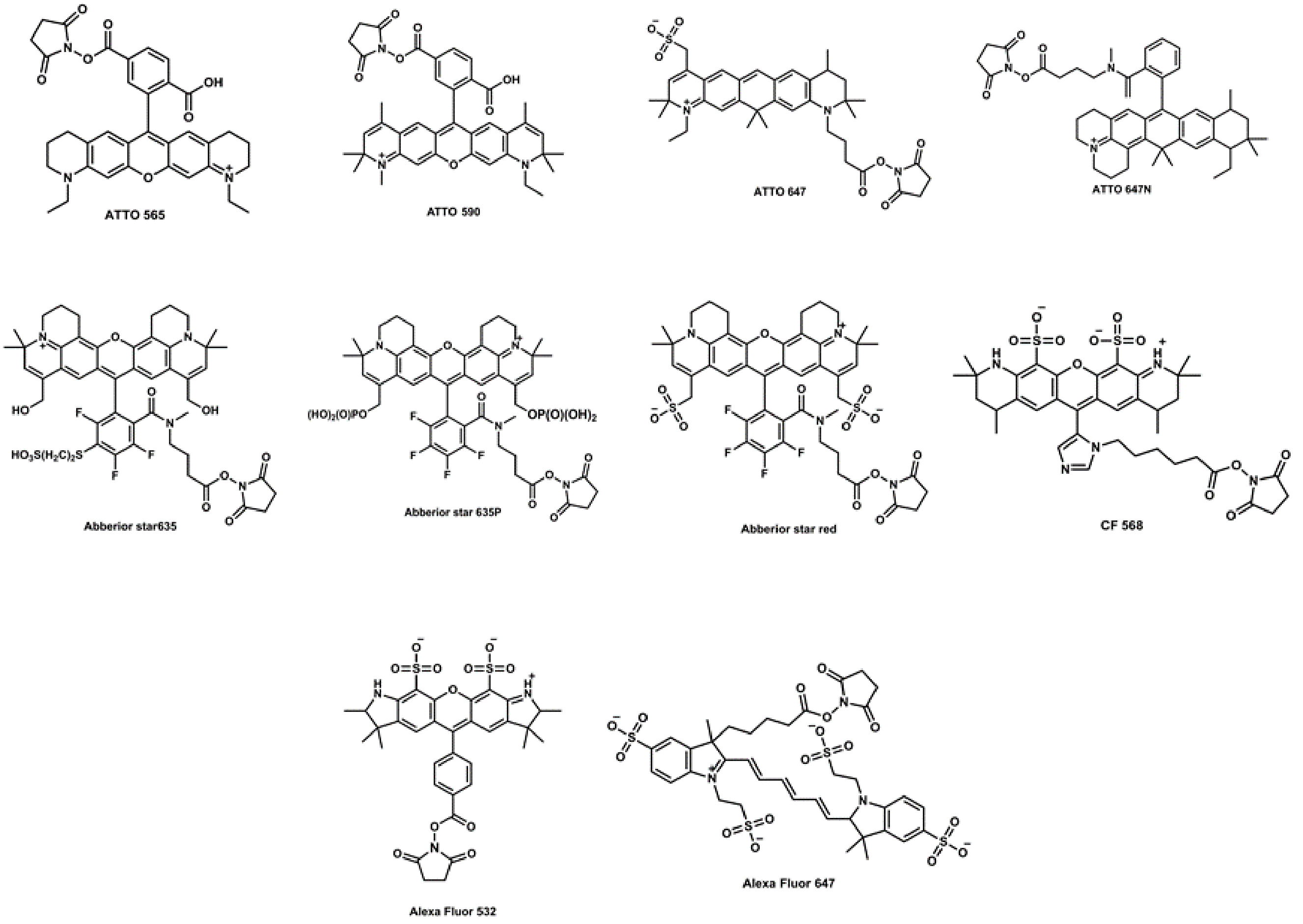

**Supplementary Fig 5:** Chemical structures of the analysed fluorophores with N-hydroxysuccinimide esters reactive group.

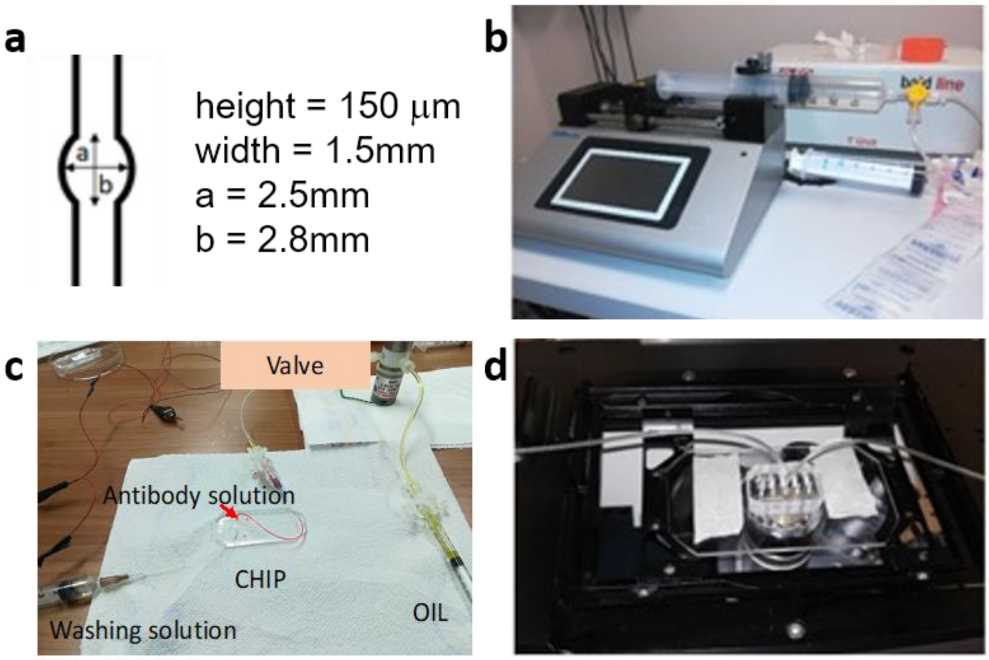

**Supplementary Fig 6: Microfluidic apparatus used for the evaluation of the relative dissociation constant for the N-FluoFab conjugates**. **A**: schematic drawing of the microfluidic chamber that was used for the serial acquisition of dye emission by confocal microscopy. **B**: injection system for the Fab-dye probes into the microfluidic chamber, consisting of, which provides a continuous flow **C**: switching system to inject simultaneously the washing buffer and the staining solution **D**: the microfluidic chamber is positioned on the stage of a confocal microscope and the flow-driven dissociation of the antigen-Fab couple is monitored as decrease in the fluorescent emission acquired by serial acquisitions. The chamber is filled with saturating concentration of staining solution, consisting of fluorescent Fab (10 μg/mL) diluted in a PBS solution of surfactant (vegetal saponin at 0,1% w/v).

### Supplementary Table

| STATISTIC ANALYSIS | FAB | Ab I | AbII |
|---|---|---|---|
| **Minimum** | 0.01225 | 0.02245 | 0.0494 |
| **Maximum** | 0.04305 | 0.06342 | 0.09337 |
| **Range** | 0.0308 | 0.04097 | 0.04397 |
| **5% Percentile** | 0.01545 | 0.02482 | 0.05521 |
| **95% Percentile** | 0.03893 | 0.05693 | 0.09154 |
| **95% CI of median** | | | |
| **Actual confidence level** | 96.7 | 96.7 | 96.7 |
| **Lower confidence limit** | 0.02533 | 0.03518 | 0.06508 |
| **Upper confidence limit** | 0.03021 | 0.04172 | 0.07111 |
| **Mean (nm)** | 0.02814 | 0.03904 | 0.06935 |
| **Std. Deviation** | 0.006576 | 0.008853 | 0.009885 |
| **Std. Error of Mean** | 0.0007352 | 0.0009898 | 0.001105 |
| **Lower 95% CI of mean** | 0.02668 | 0.03707 | 0.06715 |
| **Upper 95% CI of mean** | 0.0296 | 0.04101 | 0.07155 |

**Table 1**
